# Cytoskeleton dynamics control early events of lateral root initiation in Arabidopsis

**DOI:** 10.1101/559443

**Authors:** Amaya Vilches Barro, Dorothee Stöckle, Martha Thellmann, Paola Ruiz-Duarte, Lotte Bald, Marion Louveaux, Patrick von Born, Philipp Denninger, Tatsuaki Goh, Hidehiro Fukaki, Joop EM Vermeer, Alexis Maizel

## Abstract

How plant cells re-establish differential growth to initiate organs is poorly understood. Morphogenesis of lateral roots relies on the tightly controlled radial expansion and asymmetric division of founder cells. The cellular mechanisms that license and ensure these features are unknown. Here, we quantitatively analyse F-actin and microtubule dynamics during LR initiation. Using mutants, pharmacological and tissue-specific genetic perturbations, we show that dynamic reorganisation of both microtubule and F-actin networks is required for the asymmetric expansion of the founder cells. This cytoskeleton remodelling intertwine with auxin signalling in the pericycle and endodermis in order for founder cells to acquire a basic polarity required for initiating LR development. Our results reveal the conservation of cell remodelling and polarisation strategies between the Arabidopsis zygote and lateral root founder cells. We propose that coordinated, auxin-driven reorganisation of the cytoskeleton licenses asymmetric cell growth and divisions during embryonic and post-embryonic organogenesis.

**HIGHLIGHTS:**

- Failure for lateral root founder cells to undergo asymmetric radial expansion before division, leads to aberrant organ formation.
- Cortical microtubules arrays reorganise to facilitate this asymmetric expansion and F-actin the asymmetric division.
- Cytoskeletal reorganisation depends on auxin signalling.
- New genetic tools allow to perturb microtubules or actin in an inducible and cell-type specific manner.

## INTRODUCTION

Plants continuously form post-embryonic lateral roots (LRs) that provide anchorage and a means to forage the environment for water and nutrients. These LRs are formed from small groups of cells called founder cells which in *Arabidopsis thaliana* are recruited from the pericycle cells adjacent to xylem cells (xylem pole pericycle, XPP). LR morphogenesis is a self-organising process characterised by the regular formation of cell layers and the emergence of a typical dome-shaped primordium (Lucas et al., 2013; Malamy and Benfey, 1997; von Wangenheim et al., 2016). LR formation starts when pairs of abutting founder cells invariably swell, their nuclei migrate towards the common cell wall and an asymmetric, anticlinal division results in the formation of a stage I primordium (one cell layer). The following periclinal, formative divisions give rise to a new cell layer constituting a stage II primordium (De Rybel et al., 2010; Dubrovsky et al., 2001; Gunning et al., 1978; Laskowski et al., 1995; Malamy and Benfey, 1997). Tempering with these steps by interfering with auxin signalling in the founder cells or the overlying endodermis (Fukaki et al., 2002, 2005; Vermeer et al., 2014) blocks LR initiation, whereas altering cell wall properties (Ramakrishna et al., 2018) or the orientation of cell division planes (von Wangenheim et al., 2016) leads to formation of primordia with altered tissue organisation. The specific remodelling of the founder cells preceding their division appears therefore essential for the LR to enter the right developmental track (von Wangenheim et al., 2016).

Auxin acts as a morphogenetic trigger for LRs and is required for both their initiation and growth (Dubrovsky et al., 2008; Lavenus et al., 2013; Stoeckle et al., 2018; Vilches-Barro and Maizel, 2015). For LR initiation, auxin signalling is mediated by SOLITARY ROOT (SLR) / AUX/IAA 14 and Auxin Response Factor (ARF) proteins ARF7 as well as ARF19 (Fukaki et al., 2002; Okushima et al., 2005; Wilmoth et al., 2005). While these are necessary for founder cells to expand and enter division, the LATERAL ORGAN BOUNDARY (LBD) 16 transcription factor, target of ARF7/19, is required for their asymmetric division (Goh et al., 2012; Okushima et al., 2007). Although we have learned a lot about the hormonal control of LR formation (Bensmihen, 2015; Fukaki and Tasaka, 2009), we still lack mechanistic insights into how plants re-establish differential growth, especially in internal cell layers.

Radial expansion of LR founder cells corresponds to an increase in volume which implies generation of mechanical constraints that need to be dealt with (Vermeer et al., 2014). It has been shown that such mechanical conflict between the primordium and the overlying tissue drives the shape of the emerging LR primordium (Lucas et al., 2013). Yet, the cellular basis and mechanisms responsible for the radial expansion of LR founder cells are unknown.

Growth anisotropy in plant cells is related to the geometry of the cell and to the mechanical properties of the cell wall, in turn mainly determined by the orientation of cellulose microfibrils (Baskin et al., 1999; Landrein and Hamant, 2013; Lloyd et al., 1985). Cortical microtubules (CMTs) guide cellulose synthase complexes (Gutierrez et al., 2009; Paredez et al., 2006) to define a primary scaffold for the deposition of cellulose microfibrils (Landrein and Hamant, 2013). CMTs are therefore prime determinants of plant cell growth anisotropy. CMTs arrays orientation also influences the selection of the division plane (Rasmussen and Bellinger, 2018). Shortly before cell division, CMTs coalesce into a plane closely associated with the nucleus (Gunning and Sammut, 1990), forming the so-called pre-prophase band that ensures the robust positioning of the new cell wall (Schaefer et al., 2017). The orientation of CMTs arrays has therefore a profound influence on the direction of growth and orientation of divisions. CMTs arrays orientation is determined by the geometry of the cell (Besson and Dumais, 2011; Chakrabortty et al., 2018; Hawkins et al., 2010) but is also modulated by cues such as hormones (Bouquin et al., 2003) and mechanics (Hamant et al., 2008; Landrein and Hamant, 2013; Uyttewaal et al., 2012). The F-actin cytoskeleton has been shown to affect cell shape and division through the regulation of membrane trafficking and movement of different organelles like the nucleus (Rasmussen and Bellinger, 2018; Szymanski and Staiger, 2018; Tamura et al., 2013; Yang, 2008). Microtubules (MTs) and F-actin cooperate towards the polarisation and directional elongation of cells (Fu et al., 2005; Kimata et al., 2016; Sampathkumar et al., 2011). Exemplary is the coordinated reorganisation of F-actin and CMTs networks driving cell outgrowth to the apical region and positioning of the nucleus for the first asymmetric division of the Arabidopsis zygote (Kimata et al., 2016).

Despite the importance of founder cell remodelling for LR morphogenesis, there is no detailed study scrutinizing the role of the cytoskeleton during LR formation and how this is connected with auxin signalling. Here, we combine 4-dimensional (4D) live cell imaging, mutant and pharmacological analysis as well as cell type-specific genetic perturbation of cytoskeleton dynamics to propose a mechanistic framework for the transition from LR founder cells to a stage I primordium. We show that rearrangement of the microtubule and F-actin cytoskeleton is needed for asymmetric radial expansion, whereas polar migration of the nucleus requires the F-actin network. Finally, we show that auxin signalling in the pericycle and the endodermis is paramount for founder cells to acquire a polarity crucial for the asymmetric radial expansion of division of the founder cells. This coordinated reorganisation of the F-actin and microtubule cytoskeleton in response to auxin signalling is required to initiate the correct developmental trajectory towards a LR primordium.

## RESULTS

### Asymmetric expansion of LR founder cells coincides with nuclear migration and microtubule reorientation

Before the first asymmetric cell division (ACD) marking LR initiation, pairs of founder cells invariably expand radially and their nuclei migrate toward the common cell wall (De Rybel et al., 2010; Goh et al., 2012; Vermeer et al., 2014; von Wangenheim et al., 2016). To better characterise this process on a cellular scale, we performed high resolution live imaging of *Arabidopsis* roots expressing fluorescent markers for the nucleus and the plasma membrane in which LR initiation was synchronised by gravistimulation (Péret et al., 2012) (Figure 1A and video S1). Measurements of the cell width at the central domain, where the two founder cells are abutting, and at the peripheral domain (distal end), revealed an asymmetric radial expansion. Founder cells expand more in the central domain than at the peripheral region (Figure 1B, C and S1). This asymmetry in radial expansion is readily quantifiable before the ACD. During the nuclei rounding phase cell width in the central domain increased by 22±5% and stagnated at the periphery (3±3%; Figure 1C, average ± std. error, n=11). This asymmetric radial expansion amplifies as the primordium progresses through its development (Figure 1C). Thus, a change in founder cell shape prior to the first division prefigures the dome-shaped appearance of the LR primordium.

**Figure 1.**
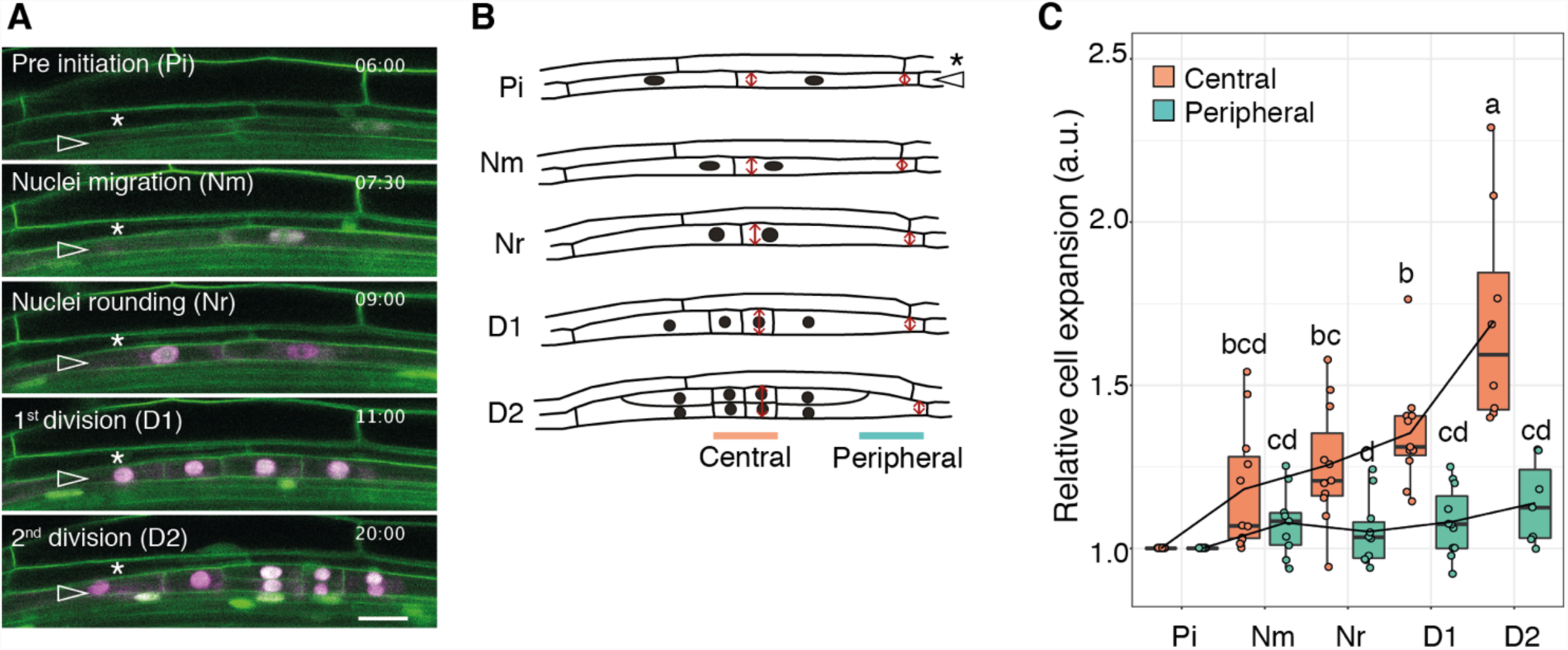
Asymmetric expansion of LR founder cells starts with nuclear migration. (A) Time-lapse image series of lateral root initiation visualised using *UB10pro::PIP1,4-3xGFP* / *GATA23pro::H2B:3xmCherry / pDR5v2pro::3xYFPnls / RPS5Apro::dtTomato:NLS* (line sC111). The time (hh:min) after plants were gravistimulated and phase of development are indicated on each panel. Images were acquired every 30 min, see also Video S1. Pre initiation, the nuclei of founder cells (open arrowhead) are positioned close to the cell centre and later migrate (Nm) towards the common cell wall and round up (Nr). The cells then divide asymmetrically (D1) and undergo a second division (D2) to form a two-layered primordium. The star indicates the endodermis. Scale bar 20 *μ*m. (B) Schematic representation of the events depicted in (A). In the central domain cells radially expand faster and prefigure the tip of the dome-shaped primordium, whereas at the peripheral domain swelling is constrained. Cell width in the central (rapidly expanding) and peripheral (not expanding) domains was measured at the position of the red arrows. The open arrowhead indicates the founder cells, the star the endodermis. (C) Quantification of founder cell expansion in the peripheral and central domains. Boxplots of normalised width for 11 founder cells in the central and peripheral domains during the indicated phases of LR initiation (see A, B). Cell width was normalised to pre initiation. Comparison between samples (n=11) was performed using ANOVA and Tukey’s HSD. Samples with identical letters do not significantly differ (α=0.05).

The CMTs arrays are prime determinants of plant cell remodelling during development (Sampathkumar et al., 2014). To visualise and quantify dynamic changes in the CMTs during the transition from founder cell to a stage I primordium, we generated *Arabidopsis* lines specifically marking MTs in the founder cells by expressing from the *GATA23* promoter (De Rybel et al., 2010) a fusion protein between green fluorescent protein (GFP) and the MICROTUBULE ASSOCIATED PROTEIN4 (MAP4) microtubule binding domain (GFP:MBD (Marc et al., 1998)). We observed that CMTs of non-dividing XPP cells are oriented in a spiral along the long axis of the cell (Figure 2A, S2). In dividing founder cells, the CMTs specifically reoriented to a more transverse orientation matching the division plane and remained in this orientation after division (Figure 2A-C, Video S2). Interestingly, as the founder cells progress through LR initiation, the CMTs arrays form distinct domains at the centre and the periphery of the primordium (Figure 2B, C). Whereas MTs in the central domain appeared isotropic in their organisation, CMTs in the peripheral domain organised in transverse parallel arrays (Figure 2C). This differential CMTs organisation is readily apparent right after the first division and reinforces in the following hours (Video S2). To quantify this differential organisation, we analysed orientation and organisation of the CMTs (Boudaoud et al., 2014; Louveaux and Boudaoud, 2018) in the central and peripheral domains of the LR founder cells before and after division (Figure 2D, E, Figure S4). Before division, CMTs arrange in anisotropic arrays at oblique to the cell long axis. After division, CMTs in the central domain tend to arrange in isotropic arrays (Figure 2D), whereas CMTs in the peripheral domain formed ordered arrays transverse to the cell long axis (Figure 2D, E, Figure S4). We observed similar dynamics of CMTs orientation using the microtubule reporter Citrine-TUA6, qualitatively corroborating our findings with the MBD marker. As the GFP:MBD reporter produced high-quality images and *GATA23pro::GFP:MBD* plants did not show compromised LR formation (Figure S3) we used this reporter to characterise CMTs in founder cells. Taken together, during the transition from founder cell to a stage I primordium, CMTs reorganise to form an isotropic array in the central area undergoing expansion and a more organized array in the area with restricted expansion.

**Figure 2.**
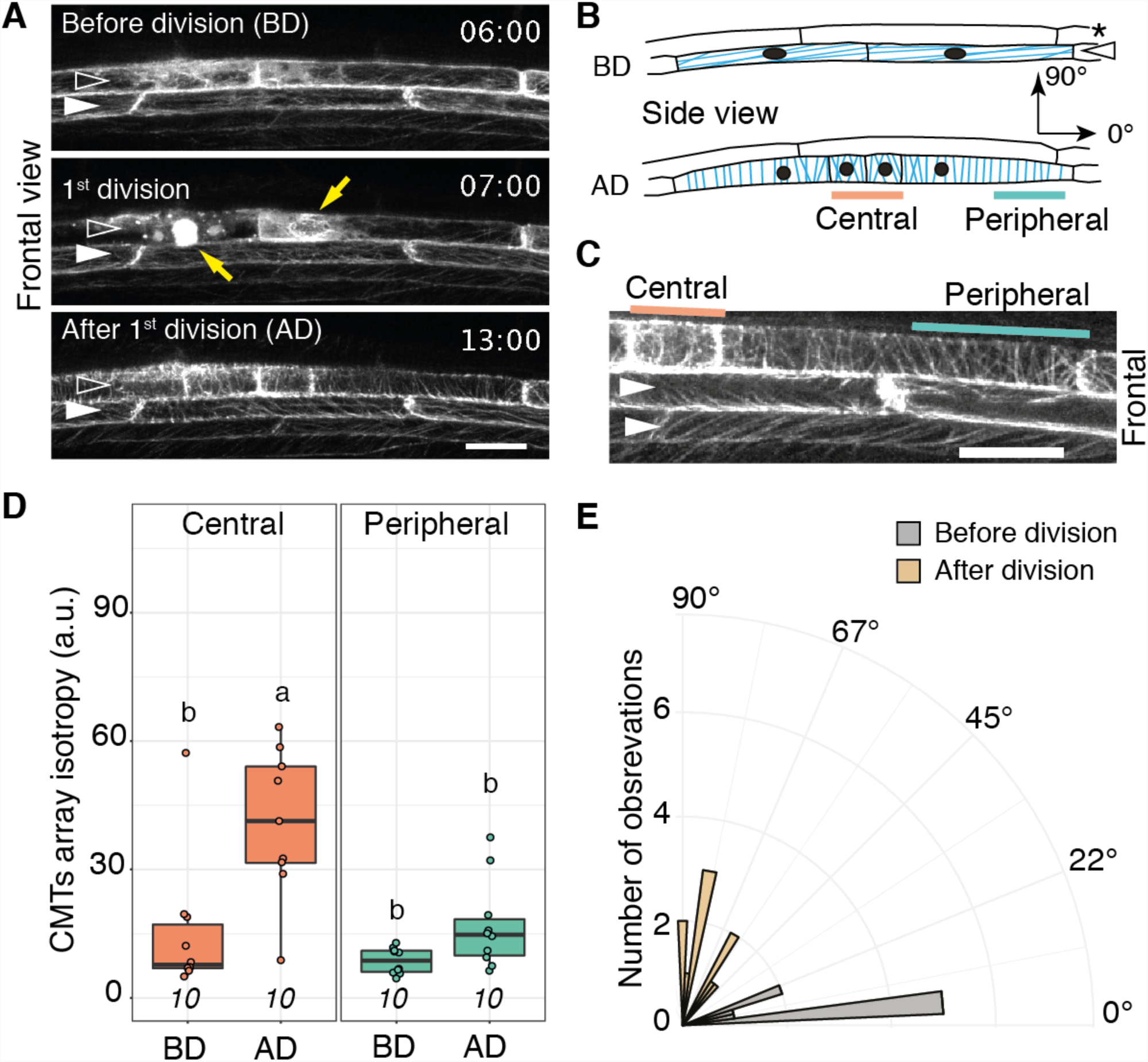
Reorganisation of microtubules in two domains during LR initiation. (A) Two-photon time-lapse image series of MTs before, during and after the first division of the founder cells visualised using GATA23pro::GFP:MBD. The time (hh:min) after plants were gravistimulated is indicated on each panel. Images were taken every 30 min, see also Video S2. Images are taken in frontal view (primordia growing toward observer, (von Wangenheim et al., 2016)) and two xylem pole pericycle (XPP) cells are visible. The open arrowheads indicate the founder cells and the filled arrowheads indicate non-dividing XPP cells. The yellow arrows indicate phragmoplast and perinuclear MTs. Scale bar 20 μm. (B) Schematic side view representation of the organisation of CMTs (blue) during LR initiation. Before division, CMTs in the XPP orient along the long axis of the cell (0°). After division, CMTs reorient by 90° to form a central, isotropic and a peripheral, anisotropic domain of CMTs organisation. The open arrowheads indicate the founder cells, the star the endodermis. (C) Closeup on CMTs in XPP cells after the first division of the founder cell (frontal view). Orange and green lines mark the zones in which orientation and isotropy of CMTs were quantified in the central and peripheral domains, respectively. The filled arrowheads indicate non-dividing XPP cells. Scale bar 20 μm. (D) Quantification of CMTs organisation in founder cells. Boxplots of the CMTs array isotropy in the central and peripheral domains of founder cells before and after the first division. The number of observations is indicated at the bottom. Comparison between samples was performed using ANOVA and Tukey’s HSD. Samples with identical letters do not significantly differ (α=0.05). (E) Orientation of CMTs arrays in the peripheral domain of founder cells. Histogram of mean CMTs array orientation (0° to 90°, see B) in the peripheral domain before and after division.

### Intact and dynamic CMTs are required for asymmetric radial expansion of founder cells

To address whether CMTs actively contribute to asymmetric radial expansion of LR founder cells, we first monitored and quantified founder cell expansion in presence of drugs that depolymerise (oryzalin) or stabilise (taxol) CMTs. We used concentrations that affected CMTs (Figure S5) while not blocking cell division (Figure 3, Videos S3 and S4). Twelve hours of oryzalin treatment resulted in increased radial expansion of both the central and peripheral domains, contrary to mock (DMSO) controls where only the central domain expanded (Figure 3A, C, E). Conversely, treatment with the CMT-stabilising taxol hampered expansion of the central domain (Figure 3A, C, E). As the tubulin-binding drugs oryzalin and taxol show opposite effects on the radial expansion, it appears that the dynamic reorganisation of CMTs facilitates asymmetric expansion of founder cell domains.

**Figure 3.**
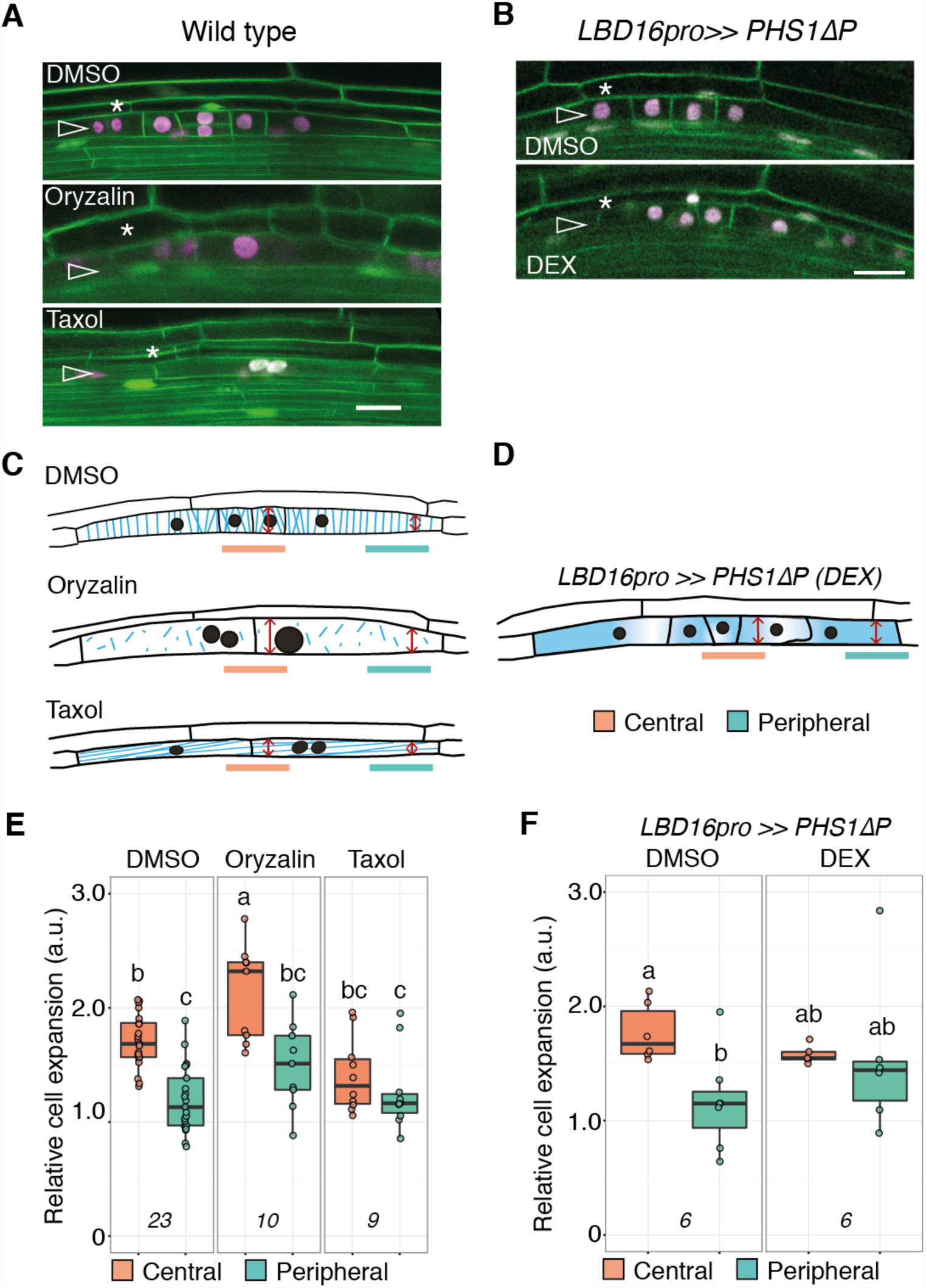
Alteration of microtubule stability and dynamics affects asymmetric radial expansion of founder cells. (A, B) Confocal sections of lateral root founder cells after the 1st division visualised using UB10pro::PIP1,4-3xGFP / GATA23pro::H2B:3xmCherry / pDR5v2pro::3xYFPnls / RPS5Apro::dtTomato:NLS (line sC111). CMTs stability or dynamics was either altered by treatment with 500 nM oryzalin or 17 μM taxol (A) or by founder cell-specific dexamethasone-inducible expression of the CMTs depolymerizing enzyme PHS1*Δ*P (B). The open arrowheads indicate the founder cells and the star the endodermis. Scale bars 20 μm. (C, D) Schematic representation of the events depicted in (A, B) with MTs in blue. Cell width in the central and peripheral domains was measured at the position of the red arrows. (E, F) Quantification of founder cell expansion. Boxplots of normalised width in the central and peripheral domains after the 1st cell division in response to oryzalin or taxol (E) or upon induction of PHS1*Δ*P in founder cells (F). Cell width was normalised to the cell width of non-dividing cells at the opposite xylem pole. The number of observations is indicated at the bottom. Comparison between samples (n_DMSO=23, n_TAXOL=10, n_ORYZALIN=9, n_DMSO=6, n_DEX=6) was performed using ANOVA and Tukey’s HSD. Samples with identical letters do not significantly differ (α=0.05).

To rule out that the effects of the drug treatments on the radial expansion of the founder cells are indirect consequences of altered CMTs in the surrounding tissues, we depolymerised CMTs in a tissue-specific and inducible manner by expressing a mutated version of the atypical tubulin kinase PROPYZAMIDE-HYPERSENSITIVE 1 (PHS1) (Fujita et al., 2013). This phosphatase-inactive PHS1 (PHS1*Δ*P) destabilised MTs, mimicking the effect of oryzalin. We created *Arabidopsis* lines expressing *PHS1ΔP* in the LR founder cells in response to dexamethasone (DEX) treatment (*LBD16pro>>PHS1ΔP*) and quantified radial expansion upon DEX or control (DMSO) treatments. Expression of *PHS1ΔP* in the *GATA23pro::MBD:GFP* background (Figure S6) verified that upon induction CMTs were indeed depolymerised in founder cells. In response to *PHS1ΔP* induction, the peripheral domain expanded more than in control conditions (Figure 3B, D, F), leading to the en bloc radial expansion of the founder cells (Figure 3I). Hence, targeted depolymerisation of CMTs in the founder cells is sufficient to observe radial expansion at the periphery.

Together, dynamic reorganisation of MTs during LR initiation defines a central domain with isotropic CMTs, permissive for radial expansion and a peripheral domain with transverse anisotropic CMTs that restrict radial expansion.

### Perturbing auxin response or founder cell polarity alters microtubule organisation and radial expansion

What contributes to the reorientation of CMTs and their organisation in two domains? Upon initiation, founder cells invariably undergo a first ACD. This division reflects the polarized nature of the founder cells and requires both cell-autonomous and non-cell-autonomous auxin signalling (Fukaki et al., 2002, 2005; Vermeer et al., 2014). We asked whether the polar organisation of CMTs and asymmetric radial expansion of founder cells is linked to the ACD and auxin signalling. To test this, we first used plants ectopically expressing *shy2-2*, a dominant repressor of auxin signalling (Tian and Reed, 1999), from the endodermis-specific *CASP1* promoter (*CASP1pro::shy2-2*). Although these plants generally fail to initiate LRs, gravistimulatedXPP cells occasionally divide (2.25±0.31 (mean ±se) divisions, n=28; Figure S7), but these divisions are symmetric and do not lead to the formation of a primordium (Vermeer et al., 2014) (Figure S7). We imaged *CASP1pro::shy2-2* seedlings, quantified CMTs orientation (Figure 4A, Figure S7, Video S5) and measured the width of XPP cells as they divided symmetrically. XPP cells did not expand radially before they divided (Figure 4D) and the CMTs did not reorganise in two domains as observed in wild type (Figure 4A). Although CMTs slightly reoriented transversally and became more isotropic, their organisation resembled still the one of non-dividing cells (Figure S7). Thus, inhibition of auxin signalling in the endodermis non-cell autonomously altered the polar organisation of CMTs and asymmetric radial expansion of founder cells in the pericycle.

**Figure 4.**
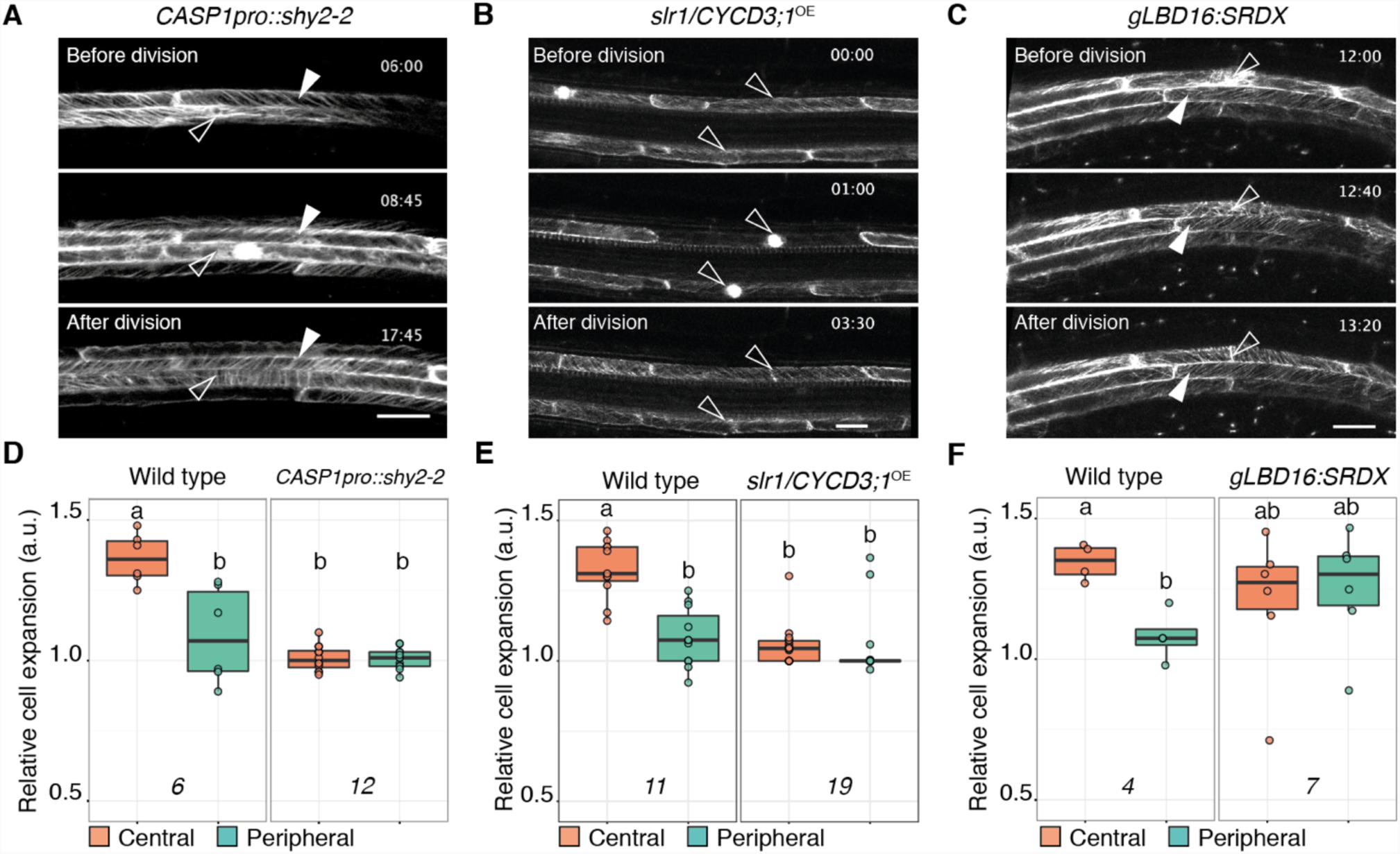
Asymmetric radial expansion of founder cells requires auxin signalling and cell polarization. (A-C) Two-photon time-lapse image series of MTs before, during and after the first division of the founder cells visualised using XPPpro::mVENUS:MBD (A) or GATA23pro::GFP:MBD (B, C) in the indicated backgrounds. The time (hh:min) after plants were gravistimulated is indicated on each panel. Images were taken every 30 min, see also Videos S5, S6 and S7. Images are taken in the frontal view and two XPP cells are visible. The open arrowheads indicate the founder cells, the filled arrowheads indicate non-dividing XPP cells. Scale bars 20 μm. (D-F) Quantification of founder cell expansion. Boxplots of normalised width in the central and peripheral domains after the 1st cell division in the indicated backgrounds. Cell width was normalised to the cell width at pre-initiation stage. The number of observations is indicated at the bottom. Comparison between samples was performed using ANOVA and Tukey’s HSD. Samples with identical letters do not significantly differ (α=0.05).

Then we tested the role of auxin signalling in the founder cells toward the polar reorganisation of CMTs and their asymmetric radial expansion. For this, we used the *slr1*/*CYCD3*^OE^ line (Vanneste et al., 2005). In this line, although auxin signalling is disrupted, the over-expression of CYCLIN D3 leads to XPP division, albeit these divisions are symmetric (Figure S8) and do not lead to formation of a primordium (Vanneste et al., 2005). We obtained very similar results to *CASP1pro::shy2-2*. Radial expansion was suppressed (Figure 4E) and CMTs arrays did not re-organise in distinct domains (Figure 4B, Figure S8, Video S6). Together, compromising auxin perception in the founder cells or in the neighbouring endodermis prevents radial expansion of founder cells and CMTs reorganisation although the cells divide. Therefore, CMTs reorganisation is not triggered by the division program and auxin signalling actively contributes to the radial expansion of founder cells and CMTs reorganisation to licence asymmetric cell expansion. To further test this, we monitored CMTs orientation in founder cells unable to polarise in response to auxin signalling. The transcription factor LBD16 is specifically expressed in the LR founder cells before the ACD (Goh et al., 2012; Okushima et al., 2007). Expression of a genomic clone of LBD16 fused to the SUPERMAN REPRESSION DOMAIN X, *gLBD16:SRDX*, a dominant repressor of LBD-mediated gene expression, blocks the polar nuclear migration in founder cells, leading to symmetric divisions (Goh et al., 2012). LR initiation stops after this division and no LRs are formed (Goh et al., 2012). We confirmed that XPP cells of *gLBD16:SRDX* plants divided symmetrically (Figure S9). Interestingly, radial expansion occurred but was comparable in the central and peripheral domains (Figure 4F), indicating that LBDs are dispensable for the radial expansion of the founder cells but required for the ACD. Quantification of CMT orientation and organisation showed that CMTs mildly reorient and form parallel arrays throughout the cell (Figure 4C, F, Figure S9, Video S7). Hence, reorganisation of CMTs in distinct domains licenses asymmetric radial expansion, requires auxin dependent cell expansion and LBD-dependent polar migration of the founder cells nuclei.

### Actin contributes to LBD16-dependent polarisation of LR founder cells

We then wondered how the polar migration of founder cell nuclei links to their radial expansion. The movement of the nucleus in plant cells depends on an intact F-actin cytoskeleton (Ketelaar et al., 2002; Tamura et al., 2013). To test whether polar migration of the nucleus is required for asymmetric radial expansion, we treated plants with the F-actin depolymerising drug latrunculin B (Lat B). Upon Lat B treatment we did not observe any coordinated migration of nuclei in founder cells, leading to symmetric divisions (Figure 5A, C, Figure S10) that resulted in formation of severely mis-formed LR primordia (Figure S10, Video S8). Although we could observe radial expansion of the founder cells, it was symmetric: the peripheral and central domains expanding similarly (Figure 5D). In these cells, CMTs organised in isotropic transverse arrays throughout the founder cell (Figure S10). To further confirm that actin-dependent nucleus migration is required for asymmetric radial expansion, we took advantage of the DeAct system, that has been developed in animal cells (Harterink et al., 2017). The expression of the mono(ADP-ribosyl)transferase domain of the *Salmonella enterica* SpvB induces ADP-ribosylation of F-actin monomers, prevents formation of filamentous F-actin and results in disassembly of all dynamic F-actin filaments (Harterink et al., 2017). We ported this system to plants (Figure S11) and generated transgenic lines that allow inducible expression of SpvB in founder cells. Upon ß-estradiol treatment, F-actin cables disappeared in founder cells (Figure S12) and their nuclei did not migrate, resulting in symmetric divisions (Figure S12). Comparable to Lat B treatment, radial expansion was symmetric (Figure 5B, C, E). Hence, polar migration of the founder cell nuclei requires the F-actin cytoskeleton and is necessary for the emergence of the two CMTs domains and the asymmetric expansion.

**Figure 5.**
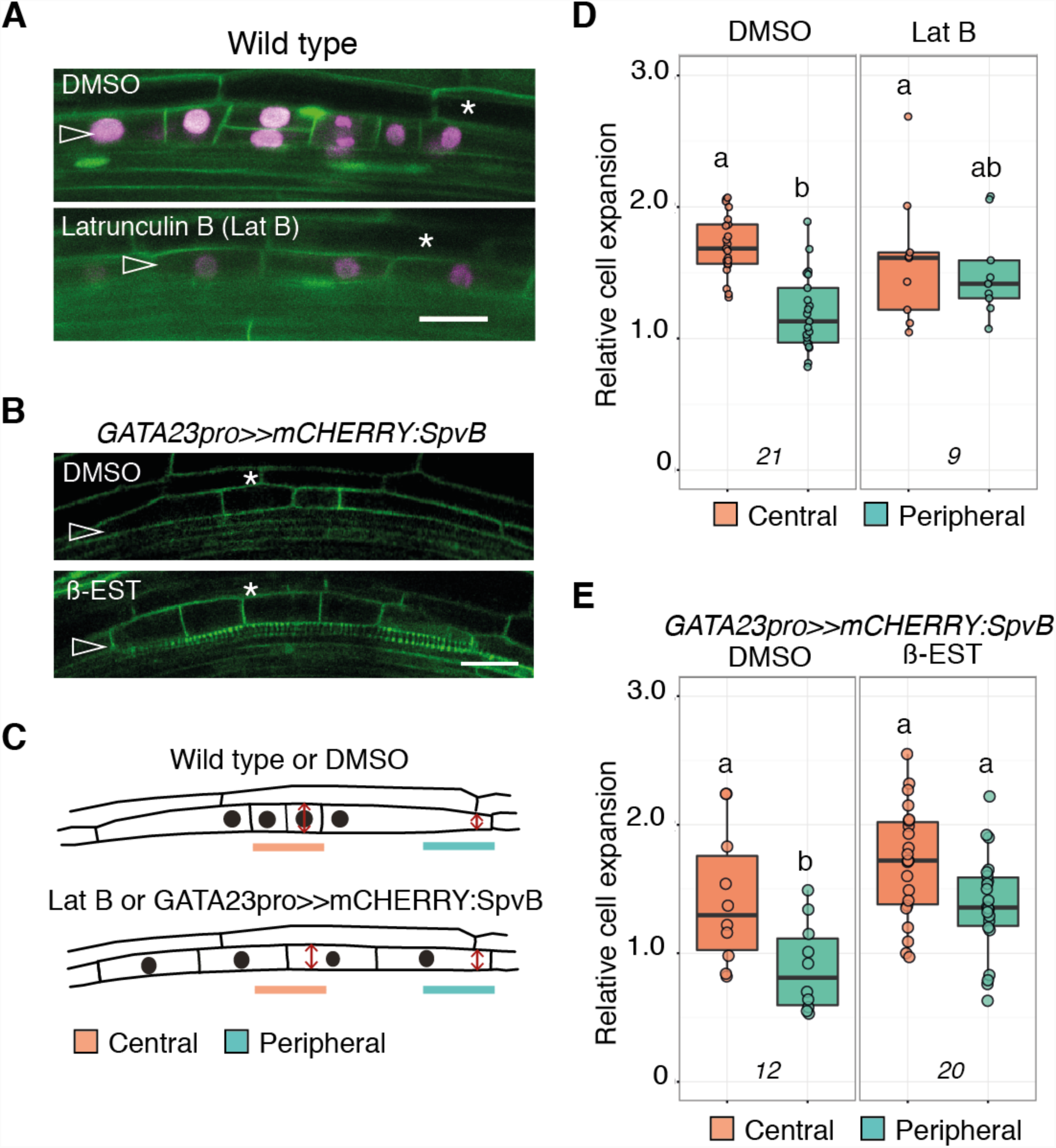
F-actin stability is required for founder cell asymmetric radial expansion. (A) Confocal sections of LRfounder cells after the 1st division visualised using *UB10pro::PIP1,4:3xGFP / GATA23pro::H2B:3xmCherry / pDR5v2pro::3xYFPnls / RPS5Apro::dtTomato:NLS* (line sC111) treated as indicated. The open arrowheads indicate the founder cells and the star the endodermis. Scale bar 20 μm. (B) Confocal sections of *GATA23pro>>mCHERRY:SpvB* LR founder cells after the 1st division visualised using *WAVE131Y* upon induction by ß-estradiol (ß-est) or control treatment (DMSO). The open arrowheads indicate the founder cells and the star the endodermis. Scale bar 20 μm. (C) Schematic representation of the events depicted in (A,B). Cell width in the central and peripheral domains was measured at the position of the red arrows. (D, E) Quantification of founder cell expansion. Boxplots of normalised width in the central and peripheral domains after the 1st cell division upon indicated treatments. Cell width was normalised to the cell width of non-dividing cells at the opposite xylem pole. The number of observations is indicated at the bottom. Comparison between samples ((A) *n_DMSO* = 21, *n_LatB* = 9; (B) *n_DMSO*=12, *n_ß-EST*=20) was performed using ANOVA and Tukey’s HSD. Samples with identical letters do not significantly differ (α=0.05).

What is the link between *LBD-*dependent auxin signalling and F-actin network reorganisation? We first set to document F-actin dynamics in founder cells of wild type plants expressing the second actin-binding domain of fimbrin tagged with GFP (ABD2:GFP) (Sano et al., 2005) driven from the *LBD16* promoter (*LBDD16pro::GFP:ABD2*). We observed F-actin bundles along the longitudinal axis of founder cells, which disappear to form a dense peri-nuclear mesh once nuclei were positioned asymmetrically in the cell (Figure 6A, C and Video S9). By quantifying the organisation of F-actin (Louveaux and Boudaoud, 2018, 2018), we confirmed that the isotropy of the F-actin network increased after cell division (Figure S13). We then investigated the dynamics of F-actin in the *gLBD16:SRDX* background where founders cells are unable to polarise in response to auxin signalling. In *gLBD16:SRDX / LBDD16pro::GFP:ABD2* roots, we observed F-actin bundles similar to wild type, although the nuclei appeared nearly immobile. Yet, we did not observe the formation of any peri-nuclear F-actin mesh in *gLBD16:SRDX / LBDD16pro::GFP:ABD2* founder cells once the cell divided. We observed bundles radiating towards the plasma membrane from the centrally positioned nucleus (Figure 6B, D, Figure S13 and Video S10). Quantification of F-actin network isotropy revealed no changes in its organisation (Figure S13) Thus, LBD_-_mediated auxin signalling contributes to LR initiation through the coordination of F-actin dynamics, nuclear migration and asymmetric radial expansion.

**Figure 6.**
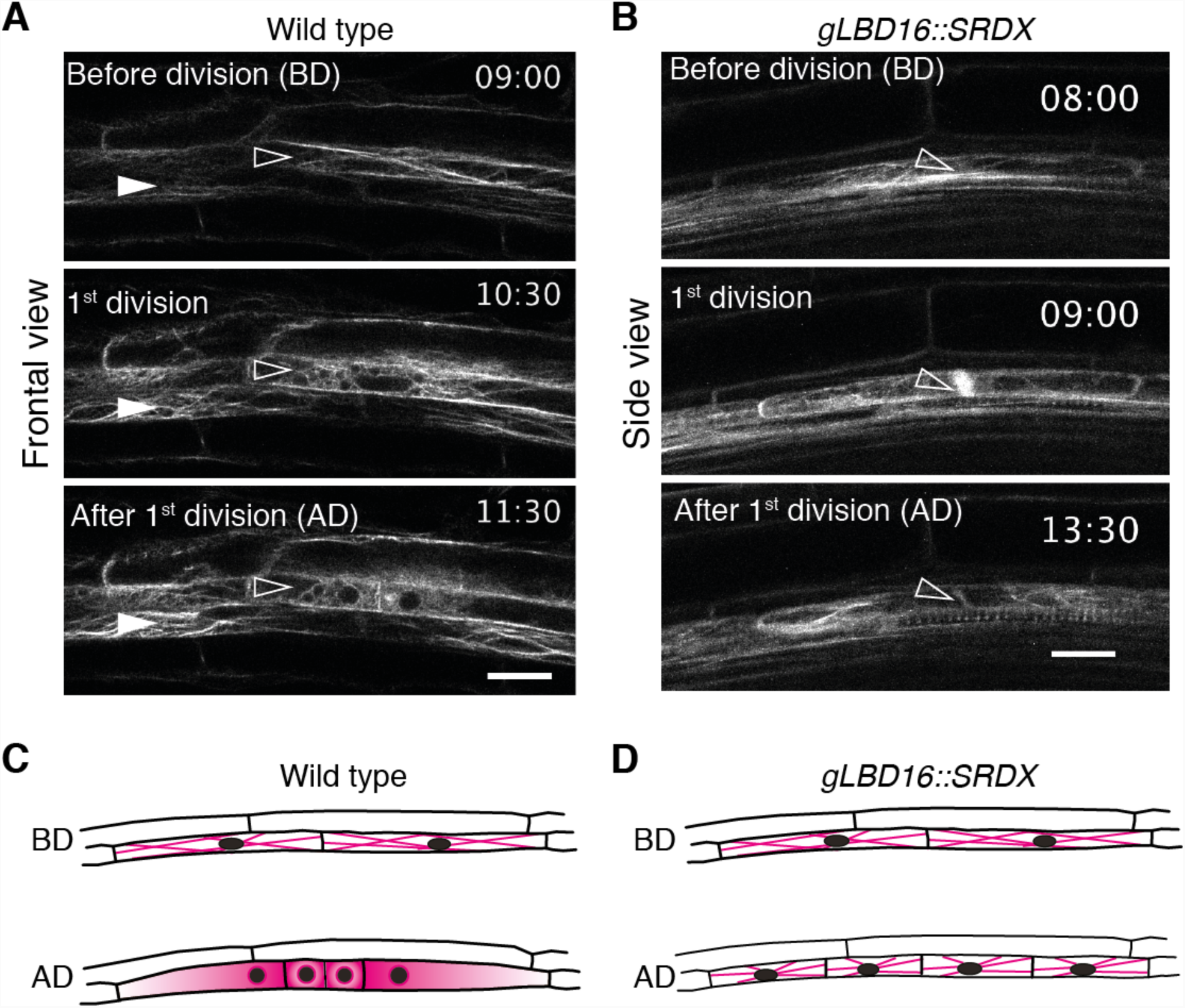
LBD16 contributes to reorganisation of F-actin network in founder cells. (A, B) Two-photon time-lapse image series of F-actin before, during and after the first division of the founder cells visualised using *LBD16pro::GFP:ABD2* in the indicated backgrounds. The time (hh:min) after plants were gravistimulated is indicated on each panel. Images were taken every 30 min, see also Videos S9 and S10. Images in (A) are taken in the frontal view, in side view in (B). The open arrowheads indicate the founder cells, the filled arrowheads indicate non-dividing XPP cells. Scale bars 20 μm. (C, D) Schematic representation of the events depicted in (A, B). F-actin is represented in magenta.

## DISCUSSION

We have precisely analysed the cellular events that precede the ACD marking the initiation of LR morphogenesis (Figure 7). We quantified the radial expansion of founder cells and observe that both founder cells coordinately and asymmetrically expand. The central domain, where the two cells are abutting, is expanding more than the distal ends, prefiguring the dome-shaped appearance of the LR primordium early on. By imaging CMTs and F-actin specifically in the pericycle and founder cells in combination with pharmacological and targeted genetic alterations, we show that F-actin and CMTs reorganisation contribute to the emergence of this asymmetric radial expansion. We observed that CMTs form two domains: isotropic arrays in the central domain and anisotropic transversal arrays at the periphery. Using mutants affected in the response to auxin signalling in the pericycle or in the overlying endodermis, we showed that auxin is important for triggering founder cell expansion and the reorganisation of CMTs arrays. Forcing cell division in the absence of auxin response, is not sufficient to reorganise CMTs. On the contrary, when founder cells can respond to auxin but are unable to polarise, they radially expand albeit symmetrically. The lack of asymmetric radial expansion is intimately linked to the polar migration of the founder cell nuclei toward the common cell wall, a process requiring the F-actin network.

**Figure 7.**
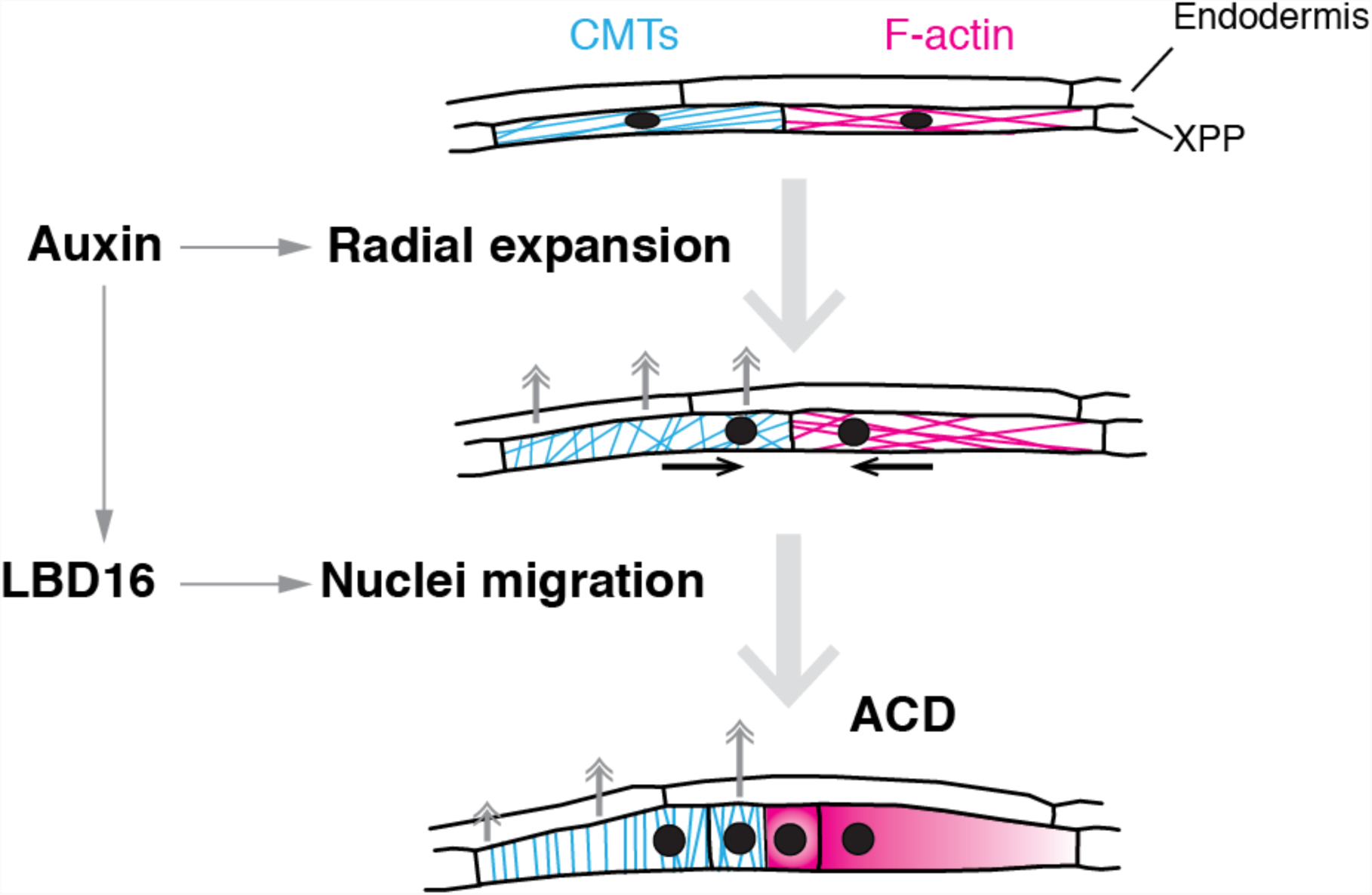
Schematic representation of the patterns and roles of CMTs and F-actin in LR initiation.

The asymmetric radial expansion of founder cells is paramount for LR morphogenesis. Modelling and empirical evidences have highlighted the importance of the proper execution of the first ACD for the subsequent morphogenesis of the LR (Ramakrishna et al., 2018; von Wangenheim et al., 2016). Here we show that when founder cells are not expanding (in *CASPpro::shy2-2* and *slr/CYCD3;1*^*OE*^), proper LR initiation does not proceed, although cell divisions can occur. When founder cells do not polarise in response to auxin (*gLBD16:SRDX*), CMT arrays are removed (*LBD16pro>>PHS1ΔP*) or their nuclei cannot migrate (Lat B treatment), they can expand radially but do it symmetrically leading to defects in subsequent LR morphogenesis (Figure S6, S10). This importance of asymmetric cell expansion for the subsequent LR development bears similarities with the first division of the *Arabidopsis* zygote. In both systems, F-actin and CMTs are coordinately remodelled to ensure the polar migration of the nucleus to the cell end undergoing isotropic expansion, setting up the stage for the zygote or founder cells to undergo ACD. This ACD gives rise to a short isodiametric daughter cell, the future embryo proper or the central domain of the LR primordium, and a long daughter cell forming the embryo suspensor or the flanks of the LR primordium. Interestingly, a transcriptomic approach has revealed that a bifurcation between the transcriptional signatures of central and peripheral domains of the LRP is established very early l(Lavenus et al., 2015), echoing the early dichotomy in cell fate observed between the daughter cells produced by the first division of the zygote (Kimata et al., 2016; Ueda et al., 2011). It is tempting to speculate that at cell scale embryonic and post-embryonic morphogenesis rely on similar mechanisms for cell remodelling and polarity acquisition. It would be interesting to follow also the vacuolar dynamics of the founder cells to see if there is a similar dynamic remodelling as described for the zygote (Kimata et al., 2019).

Beyond these parallels, LR initiation and zygote division happen in different developmental contexts and thus their cellular environment is fundamentally different. LR initiation occurs within the parental root which tissues actively accommodate the growth of the primordium (Stoeckle et al., 2018; Vilches-Barro and Maizel, 2015). The endodermis overlying the LR founder cells actively licences LR initiation via auxin signalling (Vermeer et al., 2014). It also exerts a passive role as mechanical barrier, revealed by ablation experiments. Upon removal of the endodermis, the underlying pericycle cells increase in volume and can enter division, although these divisions are not anticlinal nor asymmetric and do not lead to LR organogenesis (Marhav*ý* et al., 2016). These divisions can be partially converted to organogenic ones in presence of auxin and of dynamic CMTs (Marhav*ý* et al., 2016), reminiscent of the observations reported here. Further evidence suggests a coordinated contribution of the pericycle and the endodermis toward the radial expansion of the founder cells. Oryzalin treatment that affects CMTs not exclusively in the pericycle leads to an asymmetrical over-expansion of the founder cells (Figure 3), while the targeted removal of CMTs in the founder cells by *LBD16pro>>PHS1ΔP* lead to a symmetric radial expansion and preserve the overall magnitude. Thus, CMTs integrity in the endodermis influences the magnitude of radial expansion in founder cells. It will be interesting to precisely explore the role of CMTs in the accommodation response of the endodermis.

What drives the asymmetric expansion of founder cells? Deformation of plant cells implies modification of the balance between turgor pressure and cell wall stiffness (Landrein and Hamant, 2013). Our results show that the establishment of differential growth between the central and peripheral domains depends on the reorganisation of the CMTs. We observe that CMTs arrays in the central domain have an isotropic organisation, whereas at the periphery CMTs adopt an isotropic transversal organisation. Through their role in guiding cellulose synthase, the peripheral CMTs could reinforce the cell wall transversally and constrain radial expansion. In the central region, the shift toward an isotropic organisation would be permissive for an isotropic expansion of the cell. A shift toward isotropic CMTs organization in response to auxin signalling has been previously shown to promote cell growth and organ formation in the shoot apical meristem (Sassi et al., 2014). To our knowledge, there are no studies describing whether cellulose synthase genes show preferential accumulation or depletion in the central or peripheral domains of founder cells during the transition into a LR primordium and its subsequent development. Alternatively, this asymmetric expansion may require the local activation of EXPANSINs, proteins that can loosen the cell wall. It was recently reported that the *Arabidopsis* EXPANSIN A1 (EXPA1) is involved in the regulation of XPP cell width and the loss-of-function mutant showed aberrant cell divisions in the XPP (Cosgrove, 2000, 2016; Ramakrishna et al., 2018).

Mechanical feedback may contribute to the establishment of differential growth. It is plausible that the expanding founder cell perceives the neighbouring endodermis as an increased resistance to its expansion growth. This triggers a local reorganisation of the CMTs that in turn leads to local cell wall modification, possibly both in the founder cell and neighbouring endodermis, resulting in the asymmetric expansion. Local mechanical stimulation induces rapid, local reorganisation of the CMTs close to the stimulated area (Hardham et al., 2008) and CMTs dynamics regulated by the severing enzyme KATANIN are required to amplify differences in growth rate between adjacent cells (Uyttewaal et al., 2012).

The asymmetric expansion of founder cells depends both on auxin signalling in the XPP itself, but also in the endodermis. Interfering with either of these two components blocks the asymmetric swelling and results in an early arrest of LR development (Figure 4). Interestingly, in all cases we did not observe distinct domains of CMTs organisation. However, in the *CASP1pro::shy2-2* and *gLBD16:SRDX* backgrounds the CMTs of the XPP cells did rotate after executing a symmetric division, suggesting that a very specific process is blocked and not the capacity to change CMTs organisation, per se. In addition, in all the three auxin signalling mutants we never observed coordinated ACD, as we typically found isolated single divisions in the XPP. It appears that founder cells need to acquire a basic intrinsic polarity that is dependent on auxin-mediated signalling in both the XPP and the endodermis.

Our results highlight the intimate linkage between the polarisation of founder cells, CMTs reorganisation and asymmetric radial expansion. By manipulating the F-actin cytoskeleton we could phenocopy the loss of polarity observed with *gLBD16:SRDX* and thus showed that the F-actin network reorganisation is dependent on LBD-mediated auxin signalling. Therefore, it would be interesting to investigate which cytoskeleton-related genes are controlled by LBD16 or other members belonging to this family.

Here, we have developed two tools that allow to genetically interfere with the microtubule and F-actin cytoskeleton. To be able to control the organisation of the CMTs in a cell type-specific manner, we have introduced a genetic “oryzalin” that is based on PHS1*Δ*P (Fujita et al., 2013). This allowed us to specifically dissect the contribution of CMTs organisation in the asymmetric expansion and formation of a stage I primordium. We also generated a genetic “latrunculin”, called DeAct based on genetically encoded actin-modifying proteins (Harterink et al., 2017). Expressing SpvB in plant cells efficiently disturbed the F-actin cytoskeleton. Induction of SpvB in the founder cells interfered with asymmetric expansion, nuclear migration and resulted in symmetric divisions in the XPP. These new powerful genetic tools now provide the plant community with efficient ways to functionally probe the multifaceted roles of the plant cytoskeleton during plant development.

## AUTHORS CONTRIBUTION

AVB and DS generated resources, performed and analysed experiments. MT, PR-D, LB, ML, PVB generated resources. PD, TG and HF contributed new reagents, JEMV and AM designed, analysed the experiments and wrote the manuscript with input from all co-authors.

## ACKNOWLEDGEMENTS

We thank T. Beeckmann, N. Geldner, O. Hamant, J. Lohmann, K. Schumacher, S. Vanneste, D. Weijers for sharing published materials. We thank E. Benkova and S. Wolf for their critical reading of the manuscript. We thank the Center for Microscopy and Image Analysis of the University of Zurich for excellent technical support and assistance as well as MAPS class of 2018 for their help with the DeAct system. Work of the Maizel lab is supported by the DFG FOR2581, the Land Baden-Württemberg, the Chica und Heinz Schaller Stiftung, the CellNetworks cluster of excellence and the Boehringer Ingelheim Foundation. Work in the Vermeer lab is supported by grants from the Swiss National Science Foundation (Schweizerischer Nationalfonds zur Förderung der Wissenschaftlichen Forschung, PP00P3_157524 and 316030_164086) and the Netherlands Organization for Scientific Research (Nederlandse Organisatie voor Wetenschappelijk Onderzoek,NWO 864.13.008). DS was supported by a travel grant from the University of Zurich to visit the AM lab.

## STAR METHODS

### KEY RESOURCES TABLE

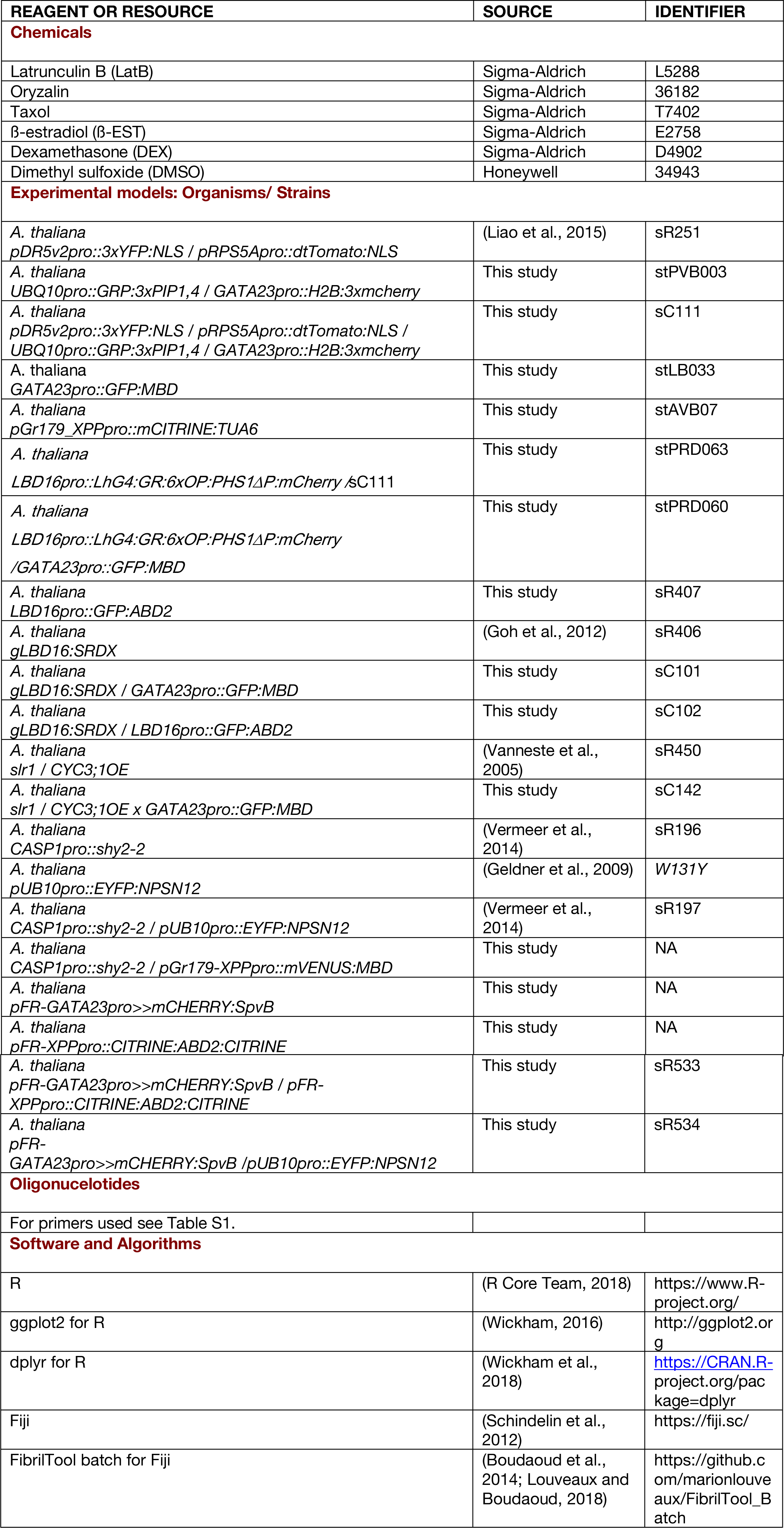

| REAGENT OR RESOURCE | SOURCE | IDENTIFIER |
| --- | --- | --- |
| **Chemicals** |  |  |
| Latrunculin B (LatB) | Sigma-Aldrich | L5288 |
| Oryzalin | Sigma-Aldrich | 36182 |
| Taxol | Sigma-Aldrich | T7402 |
| β-estradiol (β-EST) | Sigma-Aldrich | E2758 |
| Dexamethasone (DEX) | Sigma-Aldrich | D4902 |
| Dimethyl sulfoxide (DMSO) | Honeywell | 34943 |
| **Experimental models: Organisms/ Strains** |  |  |
| *A. thaliana*<br>*pDR5v2pro::3xYFP:NLS / pRPS5Apro::dtTomato:NLS* | (Liao et al., 2015) | sR251 |
| *A. thaliana*<br>*UBQ10pro::GRP:3xPIP1,4 / GATA23pro::H2B:3xmcherry* | This study | stPVB003 |
| *A. thaliana*<br>*pDR5v2pro::3xYFP:NLS / pRPS5Apro::dtTomato:NLS / UBQ10pro::GRP:3xPIP1,4 / GATA23pro::H2B:3xmcherry* | This study | sC111 |
| *A. thaliana*<br>*GATA23pro::GFP:MBD* | This study | stLB033 |
| *A. thaliana*<br>*pGr179_XPPpro::mCITRINE:TUA6* | This study | stAVB07 |
| *A. thaliana*<br>*LBD16pro::LhG4:GR:6xOP:PHS1ΔP:mCherry /sC111* | This study | stPRD063 |
| *A. thaliana*<br>*LBD16pro::LhG4:GR:6xOP:PHS1ΔP:mCherry /GATA23pro::GFP:MBD* | This study | stPRD060 |
| *A. thaliana*<br>*LBD16pro::GFP:ABD2* | This study | sR407 |
| *A. thaliana*<br>*gLBD16:SRDX* | (Goh et al., 2012) | sR406 |
| *A. thaliana*<br>*gLBD16:SRDX / GATA23pro::GFP:MBD* | This study | sC101 |
| *A. thaliana*<br>*gLBD16:SRDX / LBD16pro::GFP:ABD2* | This study | sC102 |
| *A. thaliana*<br>*slr1 / CYC3;1OE* | (Vanneste et al., 2005) | sR450 |
| *A. thaliana*<br>*slr1 / CYC3;1OE x GATA23pro::GFP:MBD* | This study | sC142 |
| *A. thaliana*<br>*CASP1pro::shy2-2* | (Vermeer et al., 2014) | sR196 |
| *A. thaliana*<br>*pUB10pro::EYFP:NPSN12* | (Geldner et al., 2009) | W131Y |
| *A. thaliana*<br>*CASP1pro::shy2-2 / pUB10pro::EYFP:NPSN12* | (Vermeer et al., 2014) | sR197 |
| *A. thaliana*<br>*CASP1pro::shy2-2 / pGr179-XPPpro::mVENUS:MBD* | This study | NA |
| *A. thaliana*<br>*pFR-GATA23pro>>mCHERRY:SpvB* | This study | NA |
| *A. thaliana*<br>*pFR-XPPpro::CITRINE:ABD2:CITRINE* | This study | NA |
| *A. thaliana*<br>*pFR-GATA23pro>>mCHERRY:SpvB / pFR-XPPpro::CITRINE:ABD2:CITRINE* | This study | sR533 |
| *A. thaliana*<br>*pFR-GATA23pro>>mCHERRY:SpvB / pUB10pro::EYFP:NPSN12* | This study | sR534 |
| **Oligonucleotides** |  |  |
| For primers used see Table S1. |  |  |
| **Software and Algorithms** |  |  |
| R | (R Core Team, 2018) | https://www.R-project.org/ |
| ggplot2 for R | (Wickham, 2016) | http://ggplot2.org |
| dplyr for R | (Wickham et al., 2018) | https://CRAN.R-project.org/package=dplyr |
| Fiji | (Schindelin et al., 2012) | https://fiji.sc/ |
| FibrilTool batch for Fiji | (Boudaoud et al., 2014; Louveaux and Boudaoud, 2018) | https://github.com/marionlouveaux/FibrilTool_Batch |

## CONTACT FOR REAGENT AND RESOURCE SHARING

Further information and requests for resources and reagents should be directed to and will be fulfilled by the lead contacts, Alexis Maizel and Joop Vermeer.

## EXPERIMENTAL MODEL AND SUBJECT DETAILS

### Plant Material and Growth Conditions

The *Arabidopsis thaliana* Columbia (Col-0) ecotype was used. Seeds were surface-sterilized (Ethanol 70% and SDS 0.1% or sodium hypochlorid 5% and 0,01% Tween 20), rinsed with ethanol 99%, air-dried and stratified at 4°C for 48h. Plants were grown vertically on 1% agar supplemented with half strength Murashige-Skoog (MS) pH5.8 at 22°C under long day or constant light. For transgene induction and pharmacological treatments, stock solutions of oryzalin, taxol, latrunculin B (Lat B), dexamethasone (DEX) and ß-estradiol (ß-EST) were dissolved in dimethyl sulfoxide (DMSO) as indicated.

The line *pDR5v2pro::3xYFP:NLS / pRPS5Apro::dtTomato:NLS / UBQ10pro::GRP:3xPIP1,4 / GATA23pro::H2B:3xmcherry* (sC111) was created by crossing sR251 (*pDR5v2pro::3xYFP:NLS / pRPS5Apro::dtTomato:NLS*) with stPVB003 (*UBQ10pro::GRP:3xPIP1,4 / GATA23pro::H2B:3xmcherry*). A F2 homozygous line was selected.

For analysis of CMTs and cell swelling in *gLBD16:SRDX*, F1 of a cross between sR406 (*gLBD16:SRDX*) and stLB033 (*GATA23pro::GFP:MBD*), were used.

For analysis of F-actin dynamics in *gLBD16:SRDX*, F1 of a cross between sR406 (*gLBD16:SRDX*) and sR407 (*LBD16pro::GFP:ABD2*), were used.

For analysis of CMTs and cell swelling in *slr1/CYC3;1*^*OE*^, F1 of a cross between sR450 (*slr1/CYC3;1*^*OE*^) and stLB033 (*GATA23pro::GFP:MBD*) were used.

For analysis of cell swelling in *CASP1pro::shy2-2* or in the DeAct system, homozygous F2 of a cross between WAVE131Y (*pUB10pro::EYFP:NPSN12*) and sR196 (*CASP1pro::shy2-2*) or *pFR-GATA23pro>>mCHERRY-SpvB* were used.

For analysis of F-actin dynamics in the DeAct system, homozygous F2 plants of a cross between *pFR-GATA23pro>>mCHERRY-SpvB* and *pFR-XPPpro::CITRINE:ABD2:CITRINE* were used.

## METHOD DETAILS

### Construction of vectors and transformation

To generate *pGr179-XPPpro::mVENUS:MBD*, we cloned the region of MAP4 as described (Dhonukshe et al., 2003) using XmaI/XbaI into pGr179-OCS3’. The XPP promoter (Andersen et al., 2018) was cloned into pGr179-MBD-OCS3’ using KpnI and mVENUS was cloned into pGr179-mVENUS:MBD-OCS3’ using EcorRI/XmaI. To generate *pGr179-XPPpro::mCITRINE:TUA6*, we cloned the XPP promoter in *pGr179-mCitrine:TUA6* (Alassimone et al., 2010) using KpnI.

To generate *GATA23pro>>mCHERRY:SpvB*, we first cloned *GATA23pro* into *p1R4-XVE* (Siligato et al., 2016) using the inFusion HD cloning kit. *pEN2-SpvB-3* was constructed with Gateway cloning using BP CLONASE II (www.thermofisher.com) after PCR amplification of SpvB (pCMV-DeAct-SpvB; Addgene #89446). Finally, all fragments were assembled in pFR-3xGW (Andersen et al., 2018) using LR CLONASE II plus (Siligato et al., 2016).

To generate *XPPpro::CITRINE:ABD2:CITRINE*, we amplified CITRINE:ABD2 (Alassimone et al., 2010) and recombined it into pEN1-2 using BP CLONASE II. All fragments we recombined into pFR-3xGW using LR CLONASE II plus.

To generate *LBD16pro::GFP:ABD2*, a fusion gene between GFP and the second actin-binding domain (ABD2) of AtFim1 was cloned into pENTR D-TOPO as described (Sano et al., 2005) then transferred into *pGWB501:LBD16pro*, which contains thee *LBD16* promoter in front of Gateway cassette (Goh et al., 2012; Nakagawa et al., 2007).

The following plasmids were generated using the Green Gate system (Lampropoulos et al., 2013). For *GATA23pro::GFP:MBD*, the following modules were combined in pGGZ001: *GATA23pro* (A), *B-dummy* (B), *GFP(A206K)* (C), *MBD* (D), *RBCS* terminator (E) and *pMAS:BastaR:tMAS* (F). The *GATA23pro* module was obtained by PCR of a 757 bp fragment corresponding to GATA23 promoter (De Rybel et al., 2010). The *MBD* module was obtained by PCR of a 1254 bp fragment of *MAP4* and removal of an internal Eco31I site by silent mutation. All other modules are described in (Lampropoulos et al., 2013).

For *UBQ10pro::GFP:3xPIP1,4 / GATA23pro::H2B:3xmCHERRY*, two intermediary modules were built: pPVB005 consisting of *UB10pro* (A), *B-dummy* (B), 3x*GFP* (C), *PIP1;4* (D), *RBCS* terminator (E) FH-adapter (F) in pGGM000 and pPVB019 consisting of *HA-adaptor, GATA23pro* (A), *H2B histone* (B), 3xmCherry (C), *D-dummy* (D), *UB10* terminator (E) and *pMAS:SulfR:t35S* (F) in pGGN000. Both modules were combined in pGGZ003 to generate the final vector. The *PIP1;4* module was obtained by PCR amplification of a 874 bp fragment of *PIP1;4* (At4g00430). All other modules are described in (Lampropoulos et al., 2013).

For *LBD16pro::LhG4:GR / 6xOP:PHS1ΔP:mCherry*, two intermediary modules were built: pLB040 consisting of *LBD16pro* (A), *B-dummy* (B), *LhG4:GR* (C), *D-dummy* (D), *RBCS* terminator (E) FH-adapter (F) in pGGM000 and pLB047 consisting of *HA-adaptor, 6xOp* (A), *B-dummy* (B), *PHS1ΔP* (C), *D-dummy* (D), *UB10* terminator (E) and *FastRed* (F) in pGGN000. Both modules were combined in pGGZ003 to generate the final vector. The *LBD16pro* module was obtained by PCR amplification of a 2157 bp fragment of *LBD16* upstream region. The *PHS1ΔP* module was obtained by PCR amplification of a 1848 bp fragment and removal of two internal Eco31I recognition by silent point mutations. The *FastRed* module was amplified from pFAST-R01 (Shimada et al., 2010) and internal Eco31I removed by silent mutations. All other modules are described in (Lampropoulos et al., 2013) and (Schürholz et al., 2018).

For transient expression in *Nicotiana benthamiana* of the DeActs system the following plasmids were built. For *UB10pro:3xGFP:SpvB*, the following modules were combined in pGGZ003: *UB10pro* (A), *B-dummy* (B), 3x*GFP* (C), *SpvB* (D), *RBCS* terminator (E) and *pMAS:BastaR:tMAS* (F). The *SpvB* module was obtained by PCR from pTetON -DeAct-SpvB (Addgene #89455). All other modules are described in (Lampropoulos et al., 2013).

For *UB10pro:3xGFP:GS1*, the following modules were combined in pGGZ003: *UB10pro* (A), *B-dummy* (B), 3x*GFP* (C), *GS1* (D), *RBCS* terminator (E) and *pMAS:BastaR:tMAS* (F). The *GS1* module was obtained by PCR from pTetON -DeAct-GS1(Addgene #89454). All other modules are described in (Lampropoulos et al., 2013).

For *UB10pro:LifeAct:-mRuby3*, the following modules were combined in pGGZ003: *UB10pro* (A), *B-dummy* (B), *LifeAct* (C), *mRuby3* (D), *HSP18.2* terminator (E) and *pUB10:HygR:tOCS* (F). The *LifeAct* module was obtained by PCR from pLifeAct-mTurquoise2 (Addgene ID: #36201), including a C-terminal GDPPVATS linker. The *mRuby3* module was obtained by PCR from pKanCMV-mClover3-mRuby3 (Addgene ID: #74252), including an N-Terminal GAGAGA linker. The HSP18.2 module was obtained by PCR on Col-0 gDNA (250 bp after STOP codon). All other modules are described in (Lampropoulos et al., 2013).

All oligonucleotides used for cloning are listed in table S1.

All plasmids were sequenced prior to usage. Arabidopsis plants were transformed using the floral dip method (Clough and Bent, 1998). At least ten individual T1 plants were analysed and one representative homozygous line was used for experiments. Tobacco leaf infiltrations were performed as described in (Moreno et al., 2013) the RNA silencing suppressor P19 was co-infiltrated (Lakatos et al., 2004).

### Microscopy

Lateral root formation was induced by gravistimulation (180° or 90°) of seedlings at 4 or 5 days after germination (DAG) and time of gravistimulation used as reference for time lapse imaging. For imaging the seedlings were transferred to a chambered cover glass as described in (Marhavy and Benkova, 2015).

Live imaging of F-actin and microtubule cytoskeleton visualised respectively with *LBD16pro::ABD2:GFP, GATA23pro::GFP:MBD* or *XPPpro::mCITRINE:TUA6* was performed on a Leica TCS SP5II confocal using two-photon (2P) equipped with a Spectra Physics MaiTai, DeepSee pulsed laser system for multi-photon excitation (GFP was excited at 800 nm). Images were taken with a 63x NA=1.30 water immersion objective. For detection a hybrid photodetector operated in photon-counting mode was used (HyD) and GFP fluorescence filtered with a 495 nm-555 nm filter. Images were taken with a resolution of 1024×1024 pixels, 400 Hz speed, maximum step size (Z-axis) of 1 μm and 3 to 4 times line averaging.

Live imaging of microtubule cytoskeleton visualised by *XPPpro::mVENUS:MBD* was performed with a Leica TCS SP8-MP, equipped with a resonant scanner (8 kHz) with either a 63x, NA=1.2 water immersion objective or a 63x, NA=1.4 oil immersion objective. Citrine and mVenus were excited with 960 nm with an insight DS+ Dual ultrafast NIR laser for multiphoton excitation. For detection, non-descanned super-sensitive photon-counting hybrid detectors (HyD), operated in photon-counting mode, were used. EYFP/mVENUS fluorescence was filtered with a CFP/YFP filter cube (483/32 & 535/30 with SP). For more information see https://www.zmb.uzh.ch/en/Instruments-and-tools/LightMicroscopes/Multiphoton/Multiphoton-Leica-TCS-SP8-MP-(Botinst).html

Live imaging of *UB10pro::PIP1,4:3xGFP / GATA23pro::H2B:3xmCherry / pDR5v2pro::3xYFPnls / RPS5Apro::dtTomato:NLS* (line sC111) was performed with a confocal microscope Leica TCS SP8 SMD (Leica microsystems). Images were taken with a HCX PL APO 63x, NA=1.30 water immersion objective. Images were taken with a resolution of 1024×1024 pixels, 400 Hz speed, maximum step size (Z-axis) of 1μm and 3 to 4 times line averaging. Fluorescent reporters were excited with an OPSLO laser (wavelengths for GFP/YFP 480 nm and 552 nm for mCherry). For detection PMT detectors were used. GFP/YFP emission fluorescence was collected in a 493-583 nm window and 583-783 nm for mCherry and dtTomato.

Imaging of tobacco leaves was performed on a Leica SPE confocal microscope. Images were taken with an 63x, NA=1.30 oil immersion objective at a resolution of 512×512 pixels. GFP as excited at 488 nm and mRuby at 532 nm. For detection PMT detectors were used. GFP emission fluorescence was collected in a 490-550 nm window and 620-670 nm for mRuby.

### Analysis of radial expansion

Quantification in wild type: sC111 seedlings were live-imaged 4h after gravistimulation every 30 min for 14 to 16 h. Five events were identified in the resulting time-lapse data: (I) pre-initiation (Pi, earliest time point, no sign of initiation), (II) nuclei migration (Nm), (III) nuclei rounding (Nr) before division, (IV) first asymmetric division (D1), (V) the central cells undergo a periclinal division (D2). For each of these events, single middle plane images were selected from the z-stacks and the width of the cell was measured at the central and peripheral domains with Fiji measure tool. Relative cell expansion in the central and peripheral domains was computed by dividing the cell width at each time point by the value at pre-initiation.

Quantification in mutants: for *CASP1pro::shy2-2, WAVE131Y* and *CASP1pro::shy2-2/WAVE131Y* seedlings (5 DAG) line were gravistimulated (90°) for 5 and 6 hours, respectively due to the developmental delay of initiation events between the two genotypes before imaging. Time-lapse z-stacks were acquired for 14 to 16 hours every 15 to 22 min. For quantification in *slr1/CYCD3;1*^*OE*^, *slr1/CYCD3;1*^*OE*^*/GATA23pro:GFP:MBD* seedlings (4 DAG) were imaged immediately shootward the differentiation zone (this line being agravitropic). Time-lapse imaging data were acquired for 14 to 16 h by recording z-stacks with intervals of 30 min. For quantification in *gLBD16:SRDX, gLBD16:SRDX/GATA23pro::MBD:GFP* seedlings (4 DAG) were gravistimulated (180°) for 7 h before imaging. Time-lapse z-stacks were acquired for 14 to 16 hours every 40 min. In all cases, analysis of cell width was performed as described for wild type, before and after the first division.

Quantification of cell width upon disruption of CMTs or actin: 4 DAG seedlings of the sC111 line were transferred to media containing 500 nM oryzalin, 17 μM taxol, 500 nM Latrunculin B or DMSO (control) for 3 h before gravistimulation (180°) for 12 h before imaging. For the LBD16>>PHS1ΔP, 4 DAG *LBD16pro::LhG4:GR/6xOP:PHS1ΔP:mCherry* seedlings were transferred to 10μM DEX or DMSO for 6 h before gravistimulation (180°) for 20 h before imaging. Seedlings were put back in the incubator and imaged again after 40 h. For the DeAct system, 4 DAG *GATA23pro>>mCHERRY:SpvB* seedlings were transferred to 5 μM ß-EST or DMSO for 24 h before gravistimulation (180°) for 12 h before imaging. In all cases, z-stacks corresponding to LR at stage I or II were acquired in the convex side of the bend and single median plane images were used to measure cell width in the central and peripheral domains using Fiji measurement tools. Similar measurements were performed on resting/non-dividing pericycle cells in the concave/opposite side of the bend and used to normalise cell width.

### Analysis of cytoskeleton organisation

Cytoskeleton organisation was analysed with the Fiji Plugin FibrilTool Batch (Boudaoud et al., 2014; Louveaux and Boudaoud, 2018). Z-stacks of marker lines decorating F-actin or MTs in LR founder cells were acquired every 30 min for 14 to 16 h. First, stacks were rotated so that the longitudinal axis of cells is horizontal (0°). Then, single planes corresponding to the cell cortex before and after cell division are identified and ROIs, excluding the cell edge, are drawn where cytoskeleton is analysed. FibrilTool returns the orientation (degrees) and quality of the cytoskeleton fibrils for each ROI (Figure S14). The quality is a measure of the anisotropy of the fibrils; a quality of 1 (highest anisotropy) corresponds to all fibrils parallel to each other (highest anisotropy). If fibrils have no directionality (lowest anisotropy), the quality will be 0. We computed the inverse of quality to represent the isotropy of the cytoskeleton before and after division. We used the absolute value of microtubule orientation (from 0° to 90°) to represent the direction of the fibrils.

### Analysis of asymmetry of cell division

Single median plane images of LRs at stage I were used to quantify the asymmetry of size between daughter cells after division. The length of each daughter cell was measured with Fiji measure tools and the ratio between the biggest and the smallest daughter computed.

### Number of cell divisions in *CASP1pro::shy2-2*

To determine the number of cell divisions in *CASP1pro::shy2-2, CASP1pro::shy2-2/ XPPpro::mVENUS:MBD* seedlings (5 DAG) were gravistimulated (90°) for 6 h before imaging. Time-lapse data were acquired every 15 – 40 min for 14 to 16 h. We counted the total number of pericycle cells dividing in the root convex side of the bend.

### QUANTIFICATION AND STATISTICAL ANALYSIS

All the statistical analyses used in this study and plotting were performed in R. The methods and p values are summarized in the figure legends.

