## Supplemental Material for "Cytoskeleton dynamics control early events of lateral root initiation in Arabidopsis"

FIGURE S1 (RELATED TO FIGURE 1) 2
FIGURE S2 (RELATED TO FIGURE 2) 3
FIGURE S3 (RELATED TO FIGURE 2) 4
FIGURE S4 (RELATED TO FIGURE 2) 5
FIGURE S5 (RELATED TO FIGURE 3) 6
FIGURE S6 (RELATED TO FIGURE 3) 7
FIGURE S7 (RELATED TO FIGURE 4) 8
FIGURE S8 (RELATED TO FIGURE 4) 9
FIGURE S9 (RELATED TO FIGURE 4) 10
FIGURE S10 (RELATED TO FIGURE 5) 11
FIGURE S11 (RELATED TO FIGURE 5) 12
FIGURE S12 (RELATED TO FIGURE 5) 13
FIGURE S13 (RELATED TO FIGURE 6) 14
FIGURE S14 (RELATED TO FIGURE 2 AND S13) 15
SUPPLEMENTAL VIDEOS 16
SUPPLEMENTAL TABLE S1 18

#### Figure S1 (related to Figure 1)

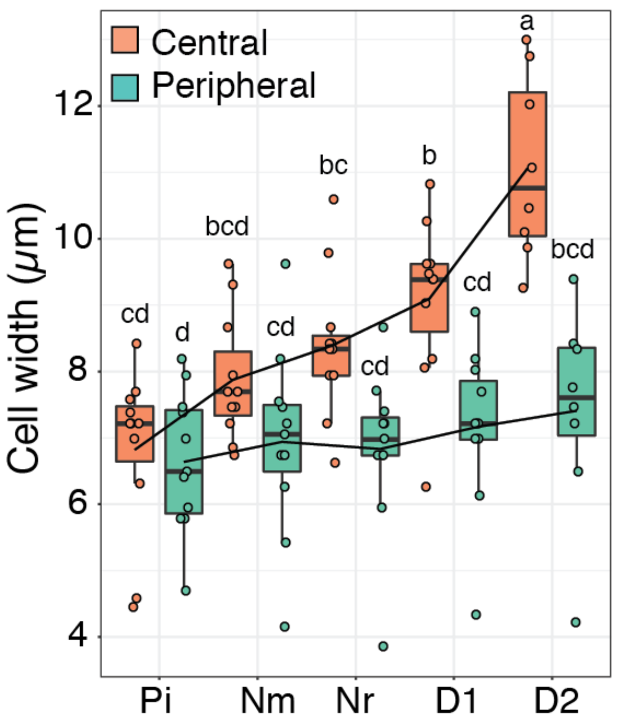


**Figure S1. Quantification of founder cell expansion in the peripheral and central domains.**

Boxplots of founder cell width in the central and peripheral domains during the indicated phases of LR initiation (see Figure 1A, B). Comparison between samples (n=11) was performed using ANOVA and Tukey's HSD. Samples with identical letters do not significantly differ ($\alpha$=0.05).

#### Figure S2 (related to Figure 2)

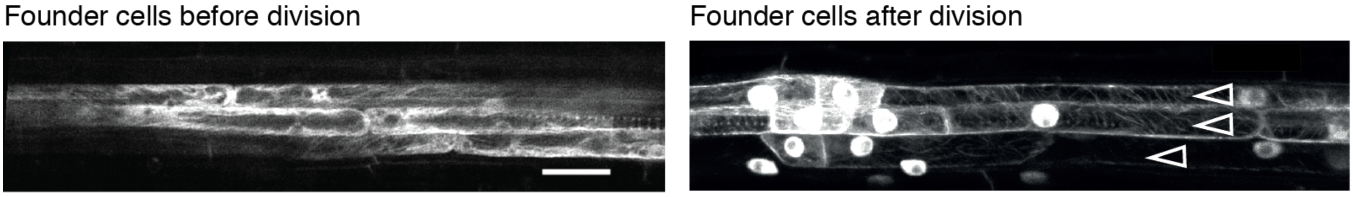


**Figure S2. CMTs organisation in pericycle cells visualised by TUA6**

Two-photon images of cortical microtubules in XPP cells and dividing founder cells visualised using *XPPpro::Citrine-TUA6* in a background expressing *GATA23pro::nls:GUS:GFP*. The open arrowheads indicate files of dividing founder cells, identified by the signal in the nucleus (*GATA23pro::nls:GUS:GFP*). Scale bar 20 μm.

#### Figure S3 (related to Figure 2)

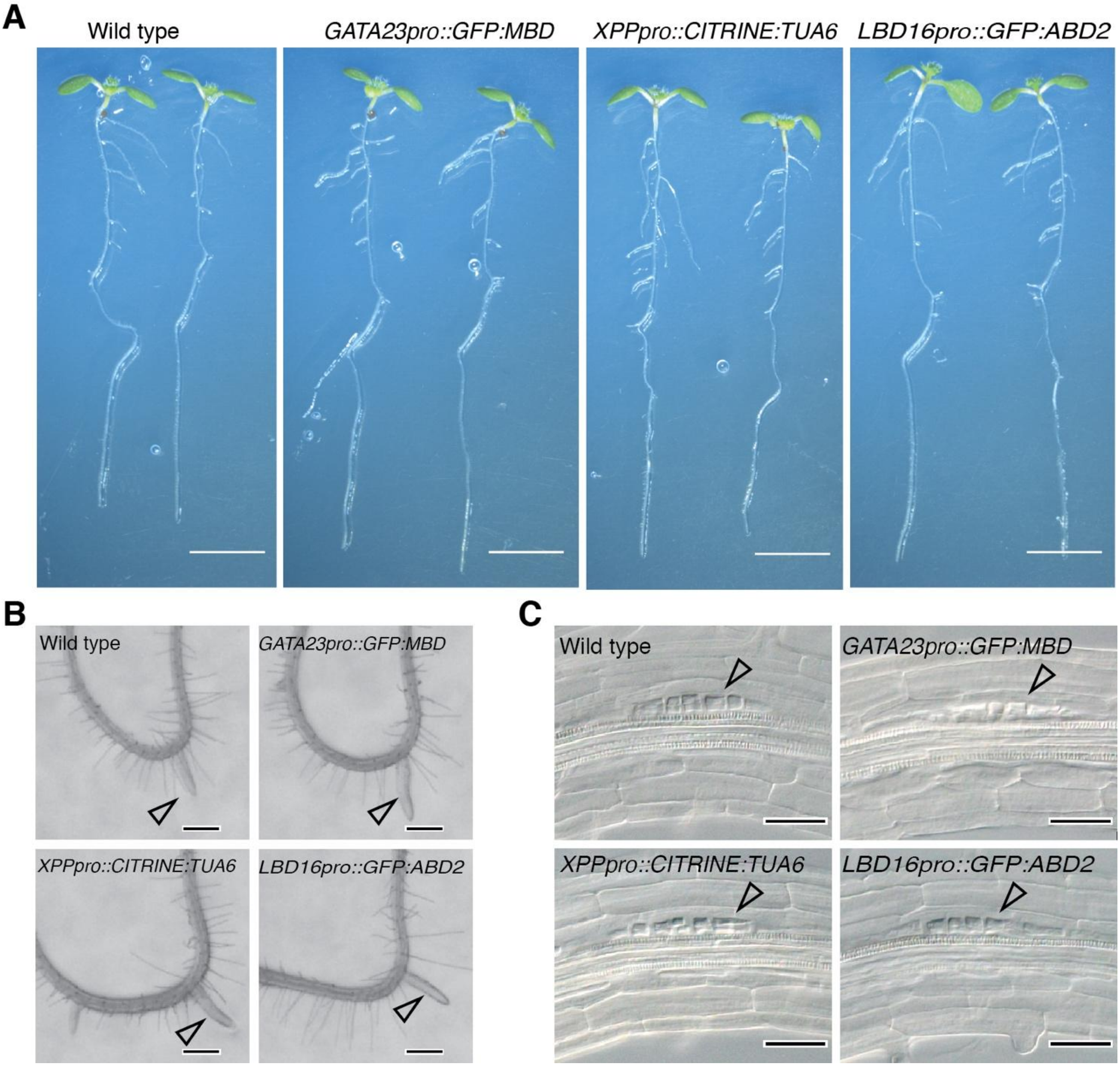


**Figure S3. Normal LR development in plants expressing markers for CMTs and actin.**

(A) Images of plants expressing the indicated cytoskeleton reporters 7 days after germination. Scale bar 500 µm. (B) Microphotographs of the gravistimulated region of roots expressing the indicated cytoskeleton reporters. Images were taken 3 days after gravistimulation, the arrowheads point to the emerged LR. Scale bars 250 µm. (C) Microphotographs of the gravistimulated region of roots expressing the indicated cytoskeleton reporters. Images were taken 24h after gravistimulation, the arrowheads point to the stage II LR primordia. Scale bars 25 µm.

#### Figure S4 (related to Figure 2)

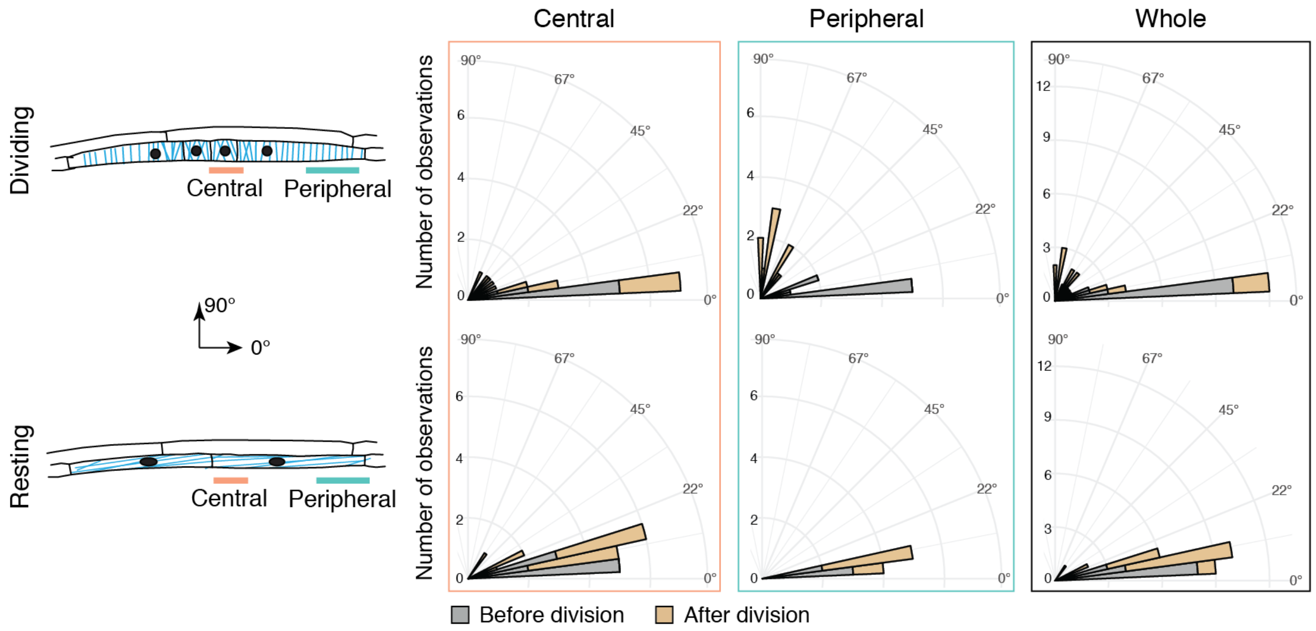


**Figure S4. Orientation of CMTs arrays in founder cells and resting XPP cells.**

Histograms of CMTs array mean orientation (0° to 90°) visualised using *GATA23pro::GFP:MBD* in the central and peripheral domains and the whole cell before and after the 1st division of founder cells. For resting (non-dividing) XPP cells, quantification "after division" were performed at the time founder cells completed division.

#### Figure S5 (related to Figure 3)

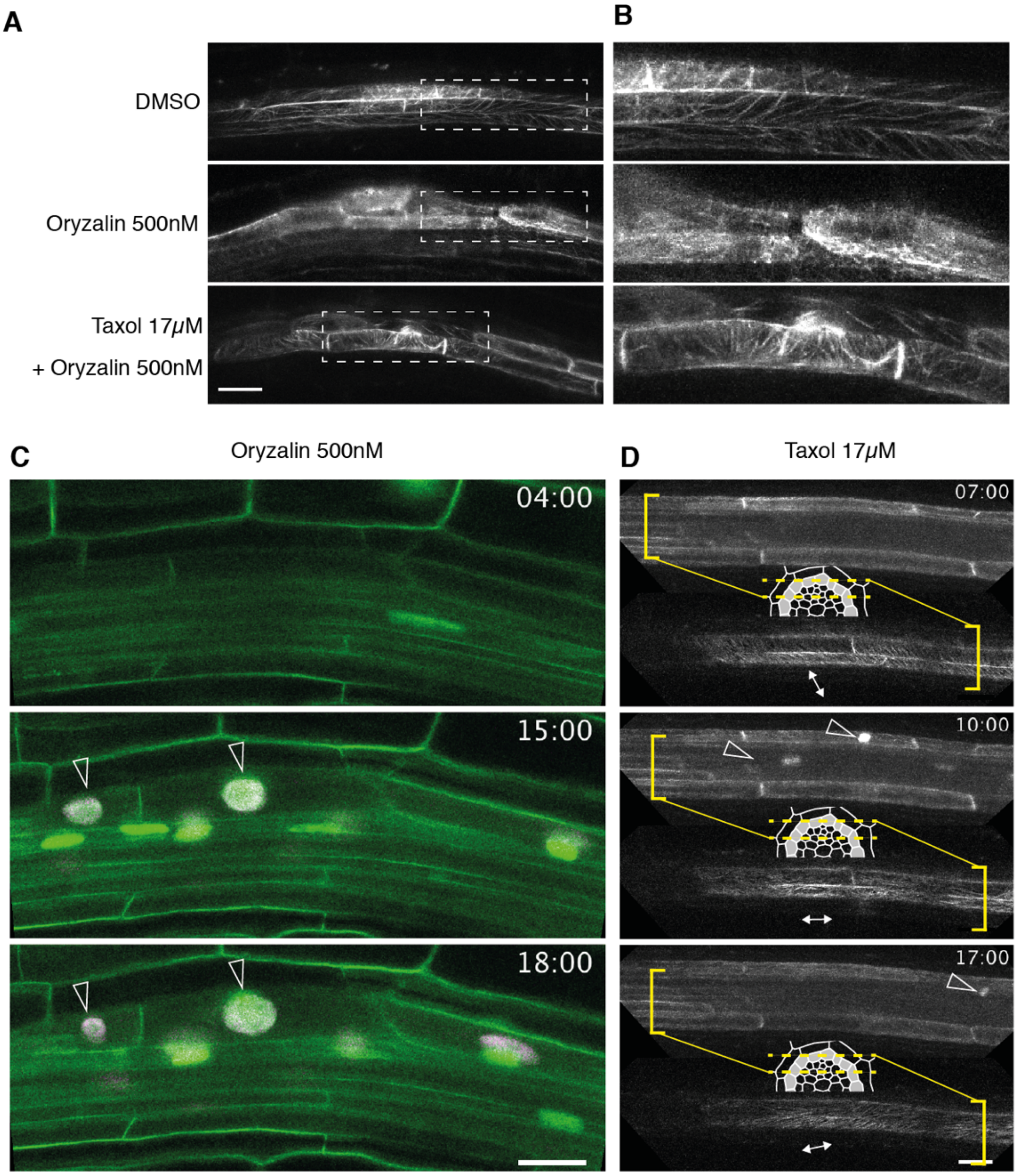


**Figure S5. Effects of oryzalin and taxol on CMTs in XPP cells**

(A, B) Two-photon images of CMTs in founder cells visualised using *GATA23pro::GFP:MBD*. (A) Plants were treated by the indicated drugs. The absence of oryzalin-induced depolymerisation of CMTs upon co-treatment with taxol indicates that the concentration of taxol used is efficiently stabilising CMTs. Scale bar 20 µm. (B) Close ups of the area boxed in (A).

(C, D) Time-lapse image series of LR initiation upon indicated treatment and visualised using *UB10pro::PIP1,4-3xGFP* / *GATA23pro::H2B:3xmCherry* / *pDR5v2pro::3xYFPnls* / *RPS5Apro::dtTomato:NLS* (C) or *GATA23pro::GFP:MBD* (D). The time (hh:min) after plants were gravistimulated is indicated on each panel and the arrowheads indicate dividing cells. Images were acquired every 30 min, see also Video S3 and S4. The diagram in D indicates the planes visualised. Scale bars 20 µm.

**Figure S6 (related to Figure 3)**

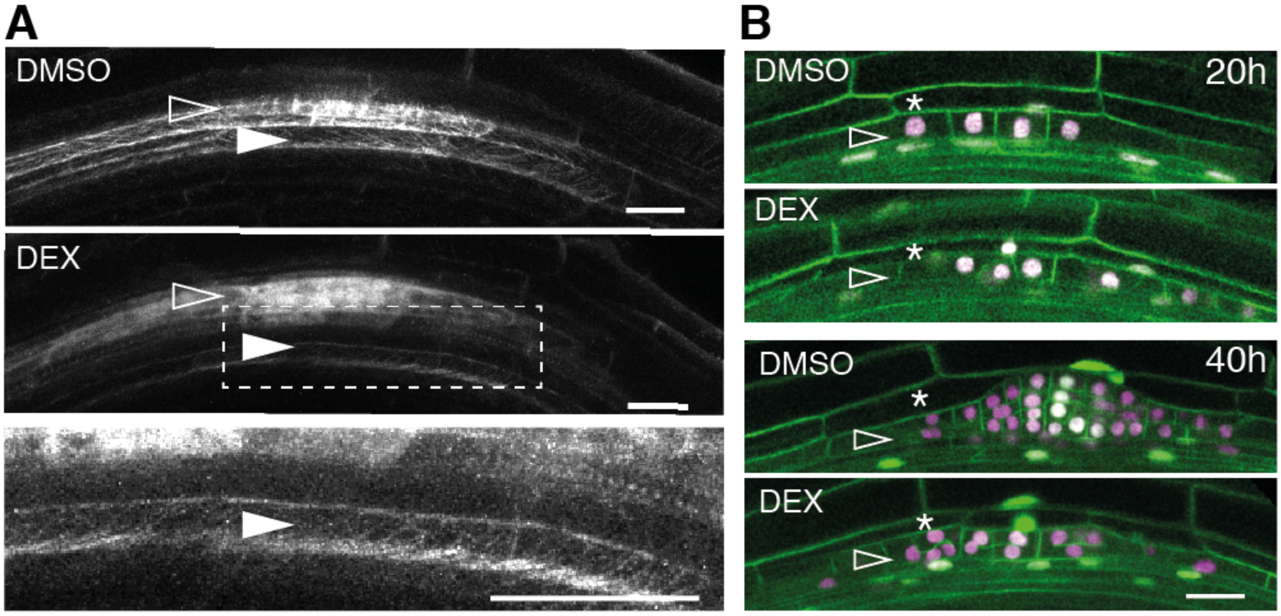


**Figure S6. PHS1ΔP-mediated perturbation of CMTs in founder cells**

(A) Two-photon images of CMTs in founder cells visualised using *GATA23pro::GFP:MBD* in the *LBD16pro>>PHS1ΔP* background. Upon *PHS1ΔP* induction (DEX) in the founder cells (open arrowheads), CMTs arrays are not visible, whereas they can be observed in non-dividing cells (closed arrowhead and inset) or in dividing founder cells in control conditions (DMSO). Scale bars 20 µm.

(B) Confocal sections of LR founder cells after the 1st division visualised using *UB10pro::PIP1,4-3xGFP* / *GATA23pro::H2B-3xmCherry* / *pDR5v2pro::3xYFPnls* / *RPS5Apro::dtTomato-NLS* (line sC111) in the *LBD16pro>>PHS1ΔP* background. Images were taken 20 or 40 h after gravistimulation upon induction of *PHS1ΔP* expression (DEX) or in control conditions (DMSO). The 20h images are the same as in Figure 3B. Founder cells are indicated by an open arrowhead, the endodermis by a star. Scale bar 20 µm.

### Figure S7 (related to Figure 4)

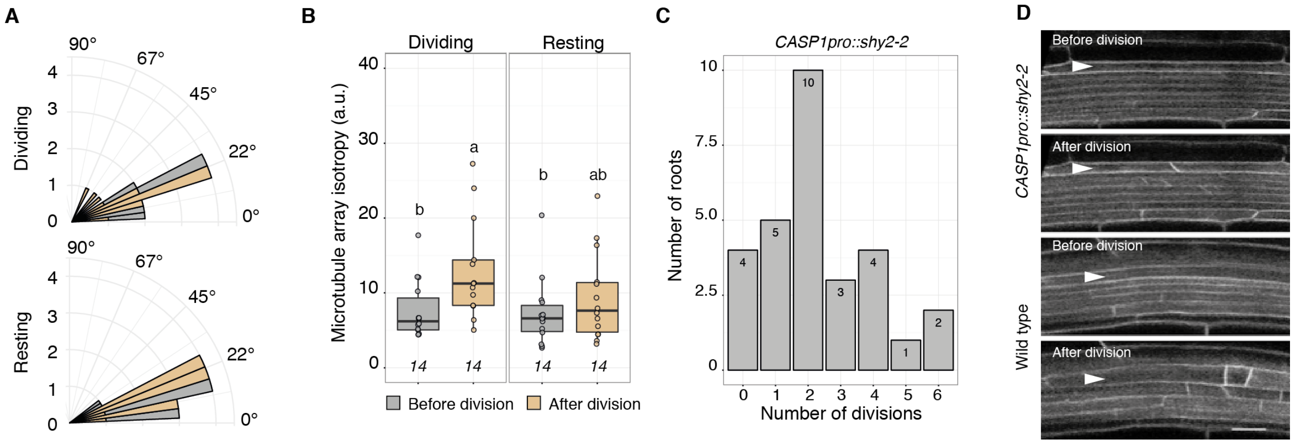


**Figure S7. CMTs array dynamics and cell division in *CASP1pro::shy2-2***

(A) Orientation of CMTs arrays. Histogram of mean CMTs array orientation (0° to 90°) in dividing and resting (non-dividing) XPP cells. CMTs orientation was identical in all parts of the cell. For resting (non-dividing) XPP cells, quantification "after division" were performed at the time founder cells completed division.

(B) Quantification of CMTs organisation in XPP cells. Boxplots of the CMTs array isotropy before and after division of XPP cells. Comparison between samples was performed using ANOVA and Tukey's HSD. The number of observations is indicated at the bottom. Samples with identical letters do not significantly differ ($\alpha$=0.05).

(C) Quantification of XPP division events. Distribution of the number of divisions of XPP cells visualised by *XPPpro:mVenus:MBD* in *CASP1pro::shy2-2* on convex side of root bend 6 h after gravistimulation.

(D) Confocal sections of LR founder cells before and after the 1st division visualised using *WAVE131Y* marker in wild type or *CASP1pro::shy2-2* backgrounds. The arrowheads indicate dividing founder cells. Scale bars 20 µm.

#### Figure S8 (related to Figure 4)

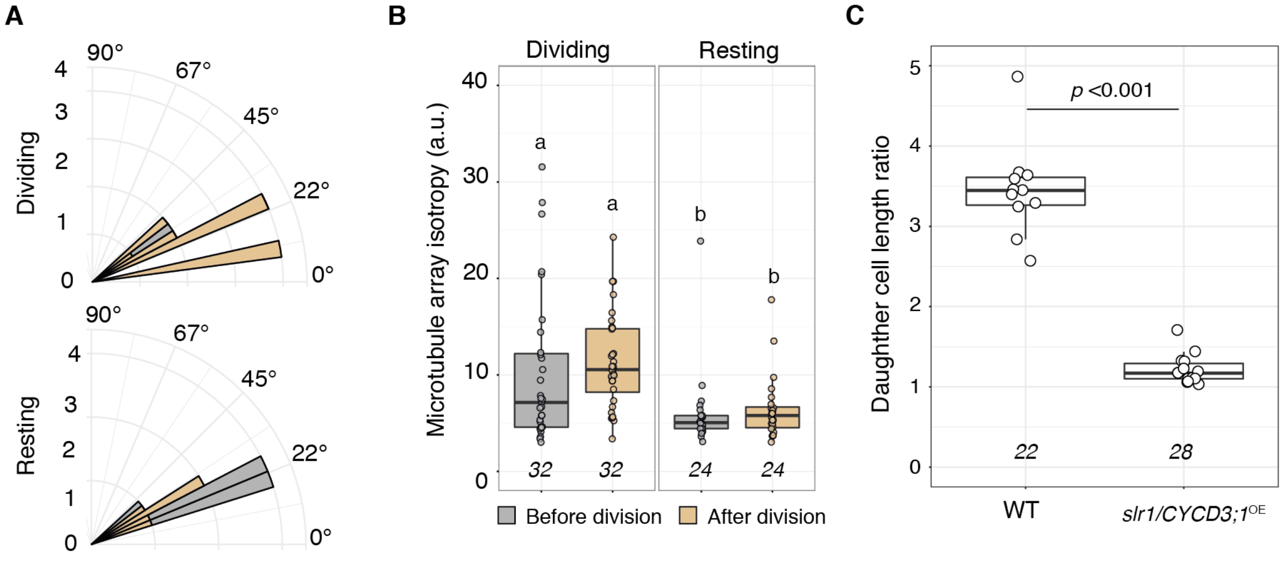


**Figure S8. CMTs array dynamics and cell division in *slr1/CYCD3;1$^{OE}$***

(A) Orientation of CMTs arrays. Histogram of mean CMTs array orientation (0° to 90°) in dividing and resting (non-dividing) XPP. CMTs orientation was identical in all parts of the cell. For resting (non-dividing) XPP cells, quantification "after division" were performed at the time founder cells completed division.

(B) Quantification of CMTs organisation in XPP cells. Boxplots of the CMTs array isotropy before and after division of XPP cells. Comparison between samples was performed using ANOVA and Tukey's HSD. The number of observations is indicated at the bottom. Samples with identical letters do not significantly differ ($\alpha$=0.05).

(C) Boxplots of the ratio of daughter cell lengths after the 1st division of founder cells in wild type (WT) and *slr1/CYCD3;1$^{OE}$*. Comparison between samples was performed using ANOVA. The number of observations is indicated at the bottom.

#### Figure S9 (related to Figure 4)

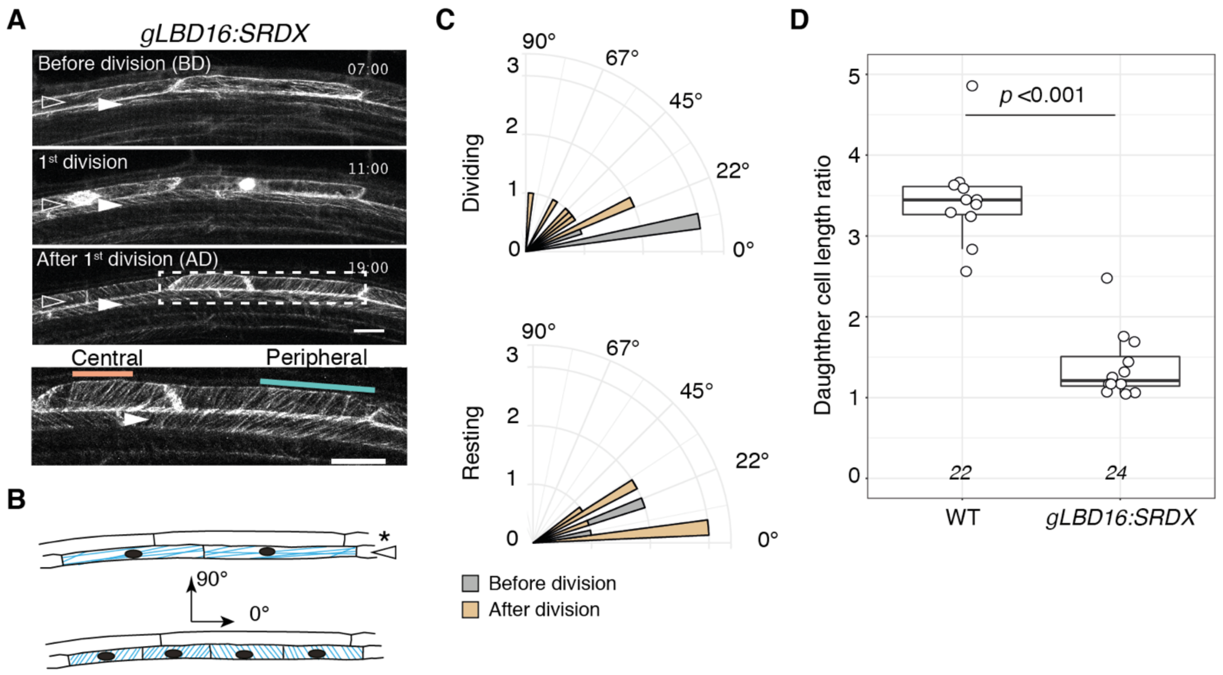


**Figure S9. CMTs array dynamics and cell division in *gLBD16:SRDX***

(A) Two-photon time-lapse image series of microtubules before, during and after the 1st division of the founder cells visualised using *GATA23pro::GFP:MBD* in the *gLBD16:SRDX* background. The time (hh:min) after plants were gravistimulated is indicated on each panel. Images were taken every 30 min, see also Video S5. Images are taken in the frontal view and two xylem pole pericycle (XPP) cells are visible. The open arrowheads indicate the founder cells, the filled arrowheads non-dividing XPP cells. Scale bars 20 μm.

(B) Schematic side view representation of the events depicted in (A). Microtubules are in blue. The open arrowheads indicate the founder cells, the star the endodermis.

(C) Orientation of CMT arrays. Histogram of mean CMTs array orientation (0° to 90°, see B) in the central and peripheral domains of dividing and resting (non-dividing) XPP cells. CMTs orientation in non-dividing cells was identical in all parts of the cell. For resting (non-dividing) XPP cells, quantification “after division” were performed at the time founder cells completed division. Vertical axis is the number of observations.

(D) Boxplots of the ratio of daughter cell lengths after the 1st division of founder cells in wild type (WT) and *gLBD16:SRDX*. The number of observations is indicated at the bottom. Comparison between samples was performed using ANOVA.

**Figure S10 (related to Figure 5)**

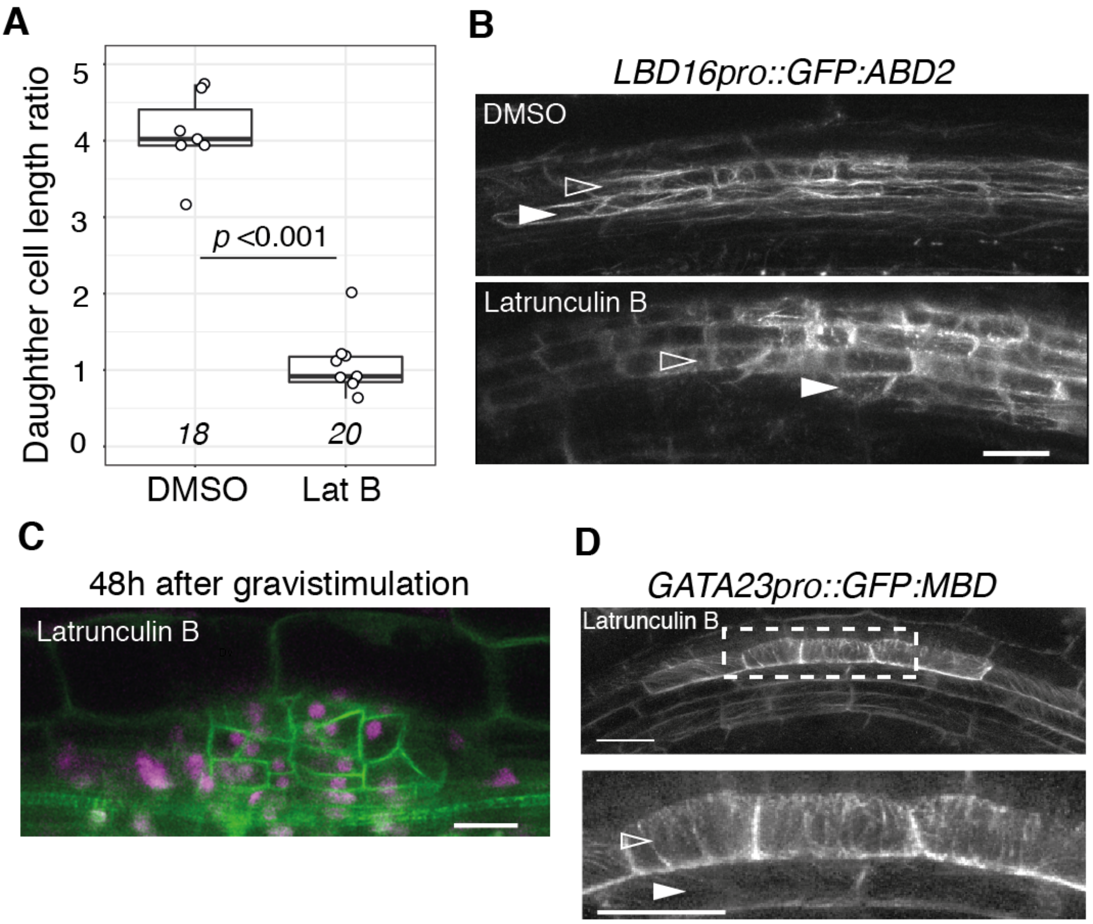


**Figure S10. Latrunculin-induced disorganization of the F-actin cytoskeleton leads to symmetric division of founder cells**

(A) Boxplots of the ratio of daughter cell lengths after the 1st division of founder cells upon latrunculin treatment. The number of observations is indicated at the bottom. Comparison between samples was performed using ANOVA.

(B) Two-photon images of the F-actin cytoskeleton upon latrunculin treatment visualised using *LBD16pro::GFP:ABD2*. The images were acquired in a frontal orientation, open arrowheads indicate the founder cells, the filled arrowheads non-dividing XPP cells. Scale bar 20 µm.

(C) Confocal section of a LR primordium 48 h after induction by gravistimulation in presence of latrunculin and visualised using *UB10pro::PIP1,4:3xGFP / GATA23pro::H2B:3xmCherry / pDR5v2pro::3xYFPnls / RPS5Apro::dtTomato:NLS* (line sC111). Scale bar 20 µm.

(D) Two-photon images of microtubules upon latrunculin treatment visualised by *GATA23pro::GFP:ABD2*. The images were acquired in a frontal orientation, open arrowheads indicate the founder cells, the filled arrowheads non-dividing XPP cells. Scale bars 20 µm.

#### Figure S11 (related to Figure 5)

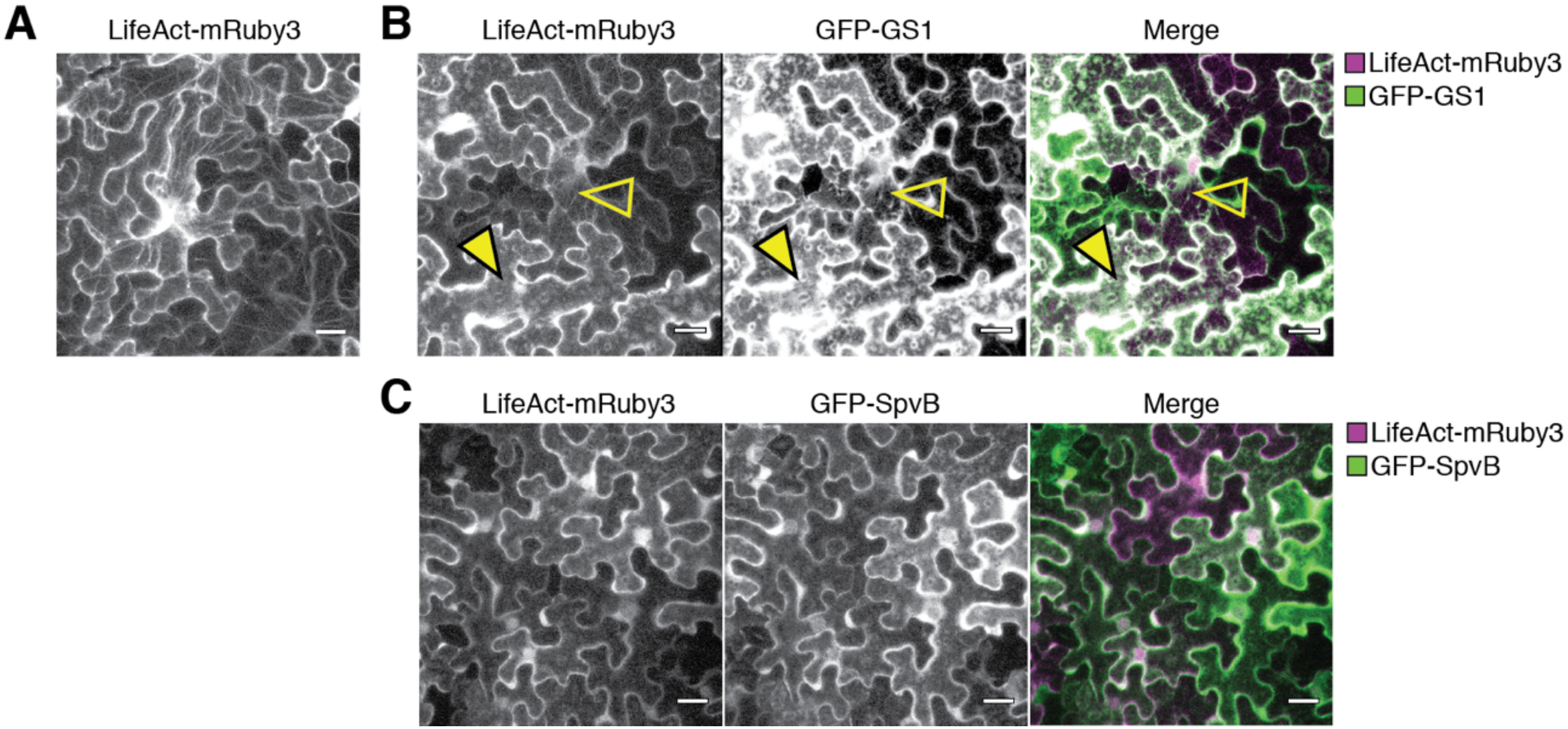


**Figure S11. The DeAct system perturbs the F-actin cytoskeleton in plant cells**

(A) F-actin visualisation in *Nicotiana benthamiana* leaf epidermis cells expressing LifeAct-mRuby3 after transient infiltration.

(B, C) Visualisation of DeAct activity by co-expression of LifeAct-mRuby3 together with either the GFP-GS1 (B) or GFP-SpvB (C). Upon expression of the stoichiometric variant (GS1), F-actin cables were only visible in cells with low levels of GS1 (B, open arrowhead) while no F-actin cable are detected in cells with higher levels of GS1 (closed arrowhead). Expression of the enzymatic DeAct variant (SpvB) leads to removal of F-actin cables in all cells (C). Scale bars 50 µm.

#### Figure S12 (related to Figure 5)

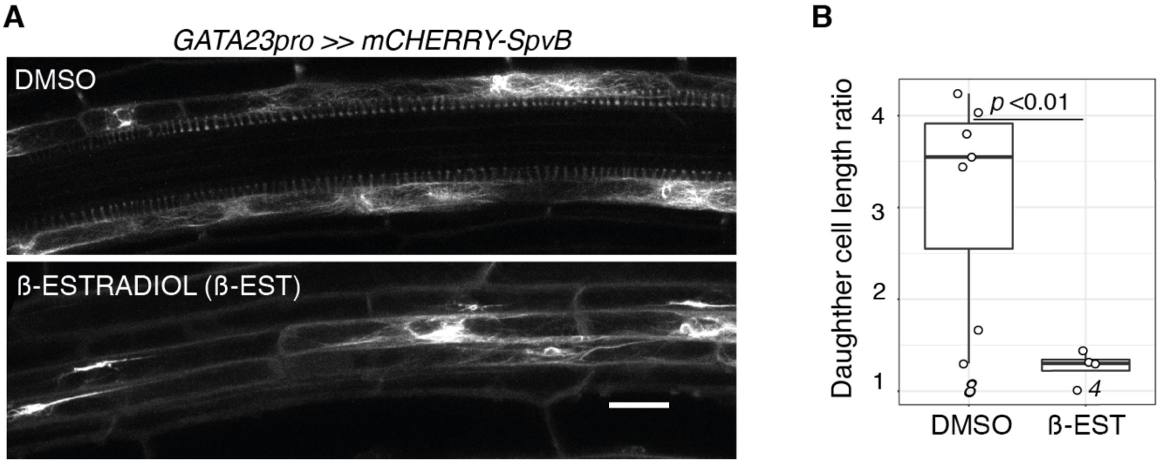


**Figure S12. The DeAct system induces disorganization of F-actin cytoskeleton and symmetric division of founder cells**

(A) Two-photon image of F-actin cytoskeleton visualised using *XPPpro::CITRINE:ABD2:CITRINE* in the *GATA23pro>>mCHERRY-SpvB* upon induction of the DeAct by ß-estradiol or in control (DMSO). Scale bars 20 µm.

(B) F-actin desorganisation leads to symmetric division. Boxplots of ratio of daughter cells length after first division of founder cells upon induction of DeAct (ß-est) or in control condition (DMSO). The number of observations is indicated at the bottom. Comparison between samples was performed using ANOVA.

#### Figure S13 (related to Figure 6)

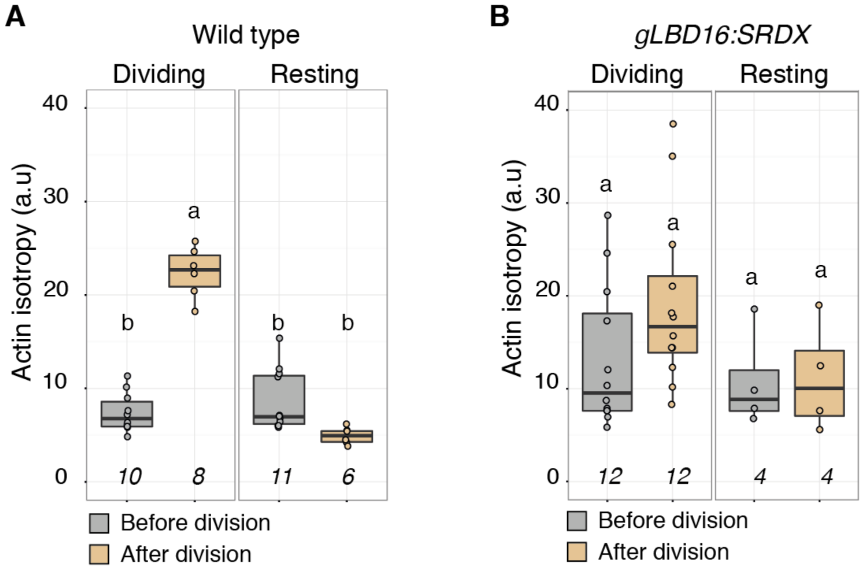


**Figure S13. Quantification of F-actin organisation in founder cells of wild type and *gLBD16:SRDX***

Boxplots of F-actin isotropy before and after division of the founder cells of wild type (A) and *gLBD16:SRDX* (B). For resting (non-dividing) XPP cells, quantification "after division" were performed at the time founder cells completed division. Comparison between samples was performed using ANOVA and Tukey's HSD. The number of observations is indicated at the bottom. Samples with identical letters do not significantly differ ($\alpha$=0.05).

#### Figure S14 (related to Figure 2 and S13)

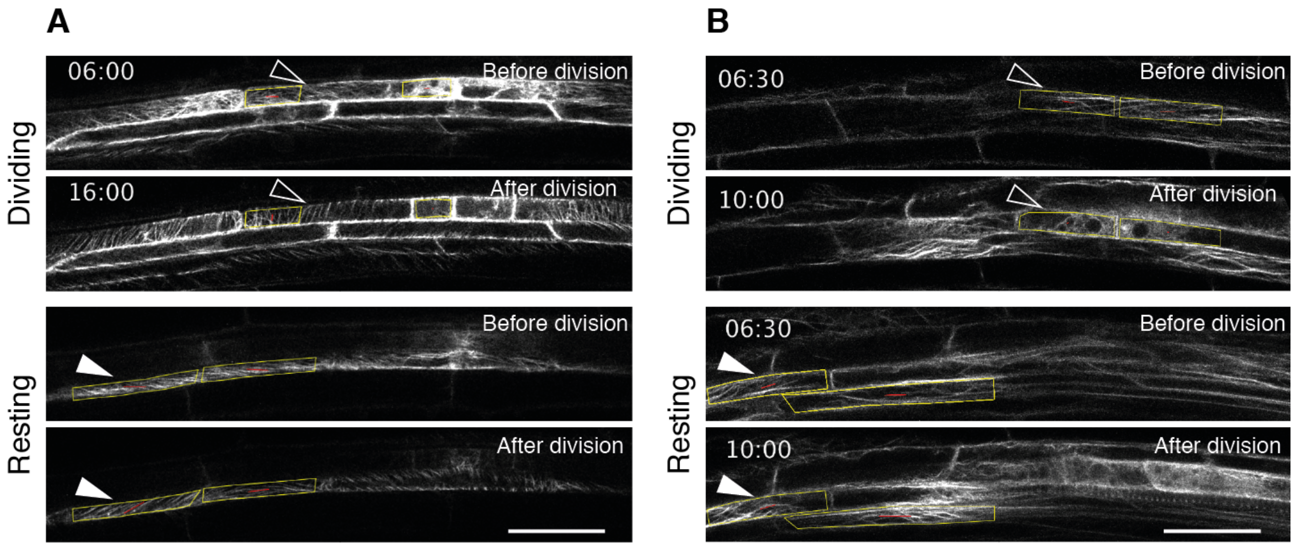


**Figure S14. Example of quantification of cytoskeleton organisation with FibrilTool**

(A, B) Two-photon image of CMTs (A) or F-actin (B) cytoskeleton visualised using *GATA23pro::GFP:MBD* (A) or *XPPpro::CITRINE:ABD2:CITRINE* (B). Images of founder cells (dividing, open arrowheads) and resting (non-dividing) XPP cells (arrowheads) taken before and after founder cell division are shown. The yellow boxes represent the ROI defined for quantification with FibrilTool. The red line within, represents the main orientation (angle to horizontal) and quality (length of line) of the fibrils as computed by FibrilTool. The time (hh:min) after plants were gravistimulated is indicated on each panel. Scale bars 20 µm.

#### Supplemental Videos

| | | |
|---|---|---|
| 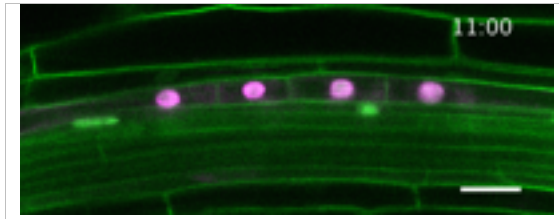  | **Video S1 (related to Figure 1)**<br>Time lapse of radial expansion in wild type | https://youtu.be/0Ike2uh6b6w |
| 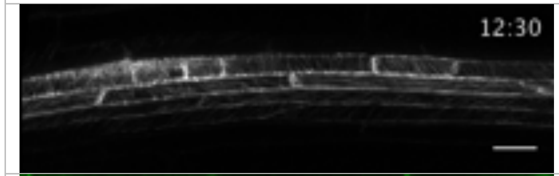  | **Video S2 (related to Figure 2)**<br>Two-photon time lapse of CMTs in founder cells in WT | https://youtu.be/WNnNmUAKuSY |
| 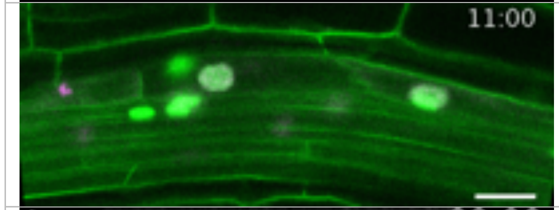  | **Video S3 (related to Figure 3)**<br>Time lapse of radial expansion in wild type upon oryzalin treatment | https://youtu.be/a2qx92Xzom0 |
| 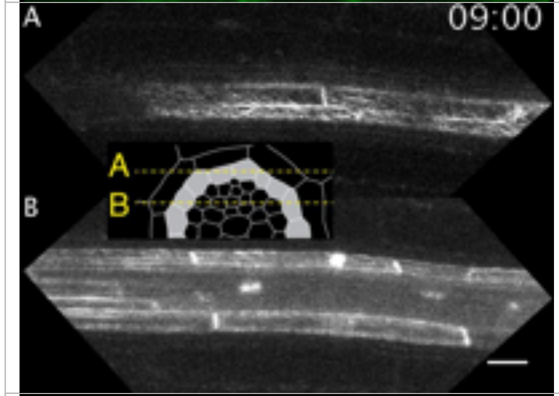  | **Video S4 (related to Figure 3)**<br>Time lapse of CMTs in founder cells of wild type upon taxol treatment | https://youtu.be/17yOWEbr06E |
| 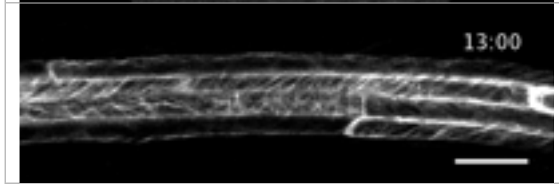  | **Video S5 (related to Figure 4)**<br>Two-photon time lapse of CMTs in founder cells in *CASP1pro::shy2-2* | https://youtu.be/vj2xZ-0vOqQ |

| | | |
|---|---|---|
| 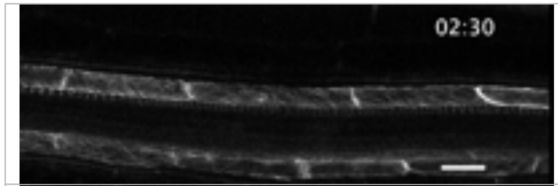 | **Video S6 (related to Figure 4)**<br>Two-photon time lapse of CMTs in founder cells in *slr/CYCD3oe* | https://youtu.be/h42KPe4kr8o |
| 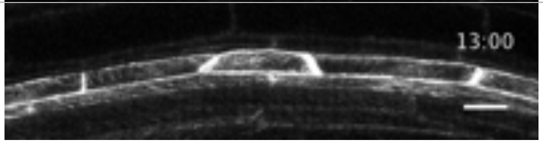 | **Video S7 (related to Figure 4)**<br>Two-photon time lapse of CMTs in founder cells in *gLBD16:SRDX* | https://youtu.be/7UW7WZ76Xp4 |
| 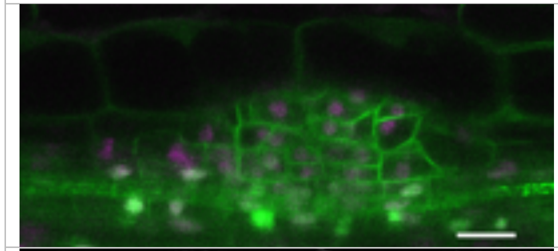 | **Video S8 (related to Figure 5 and S10)**<br>Z-stack of wild type 48h after treatment with LatB. | https://youtu.be/cuM_i4AXHAw |
| 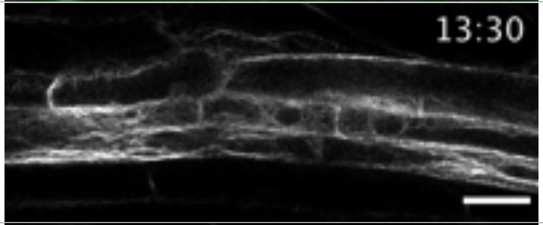 | **Video S9 (related to Figure 6)**<br>Two-photon time lapse of F-actin in founder cells in wild type | https://youtu.be/8MzprcP_cuM |
| 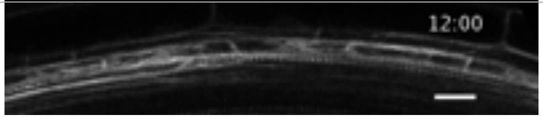 | **Video S10 (related to Figure 6)**<br>Two-photon time lapse of F-actin in founder cells in *gLBD16:SRDX* | https://youtu.be/20U4HBsHbkw |

#### Supplemental Table S1

| Purpose | Primer | Sequence |
|---|---|---|
| **XPPpro amplification** | proXPPKpnIfw | GGGGTACCGTGTTTGGTTCGTAATTA |
| | proXPPKpnIrv | GGGGTACCTTTGGAAATCTTCGTGTG |
| **MAP4 MBD amplification GW** | MAP4-XmaIfw | CCCCCCGGGGCCGACCTCAGTCTTGTG |
| | MAP4-XbaIrv | TGCTCTAGATTAGATGCTTGTCTCCTGG |
| **Venus cloning** | FP_EcoRIfw | GGAATTCATGGTGAGCAAGGGCGAG |
| | FPXmaIRV | CCCCCCGGGCTTGTACAGCTCGTCCATG |
| **SpvB amplification GW** | attB2r-SvpBfw | TGTACAAAGTGGTTGGAGGTAATTCATCTCGAC |
| | attB3-SvpBrv | ATAATAAAGTTGTTCATGAGTTGAGTACCCTC |
| **GATA23pro tet-system** | iGATA23fw | GCGCCACGATACCGGTGGTACCCATAACTTTTCAATAATGG |
| | iGATA23rv | GAATTGGTACGTACTCGAGGGTACCCAAATAAAAAAAAACAATC |
| **ABD2 cloning** | attB1FPfw | GGGGACAAGTTTGTACAAAAAAGCAGGCTTAACAatggtgagcaagggcga |
| | attB2-ABD2rv | agaaagctgggtTTTCGATGGATGCTTCCTCTG |
| ***GATA23pro* GreenGate module** | P-0976 | AACAGGTCTCAACCTATAACTTTTCAATAATGGATCTCG |
| | P-0977 | AACAGGTCTCTTGTTGAGTCATCAAGAAAGGCTTAAG |
| ***MBD* GreenGate module** | P-1601 | ACAGGTCTCATCAGGTTCCCGGCAAGAAGAAG |
| | P-1602 | ACAGGTCTCGATACCTTGGATATGTCCACTTTC |
| | P-1603 | AACAGGTCTCGGTATCCTCCAAGTGTGG |
| | P-1604 | ACAGGTCTCTGCAGTTAACCTCCTGCAGGAAAGTG |
| ***PIP1;4* GreenGate module** | P-1225 | AACAGGTCTCAGGCTCAACAATGGAAGGCAAAGAAGAAGAT |
| | P-1226 | AACAGGTCTCTCTGAACTCTTGCTCTTGAAAGGAATC |
| **SpvB GreenGate module** | P-1944 | AACAGGTCTCATCAGGTATGGGAGGTAATTCATC |
| | P-1945 | AACAGGTCTCTGCAGTTATCATGAGTTGAGTAC |
| **GS1 GreenGate module** | P-1942 | AACAGGTCTCATCAGGTATGGTGGTGGAACAC |
| | P-1943 | AACAGGTCTCTGCAGTTATCAGAATCCTGATGC |
| ***LBD16pro* GreenGate module** | P-1820 | AACAGGTCTCAACCTTACCATGAACTACAATG |
| | P-1821 | AACAGGTCTCTTGTTCGGCGAAACGAACAAA |

| | | |
|---|---|---|
| ***PHS1ΔP* GreenGate module** | P-1581 | ACAGGTCTCAGGCTCAACAATGGTCACTAGTGCAG |
| | P-1586 | ACAGGTCTCTCTGAATTAGCAGCTTTGCTAATCAATG |
| | P-1816 | CCTGTGAGACTAACATTTGATC |
| | P-1817 | GATCAAATGTTAGTCTCACAGG |
| | P-1818 | CAAATCCCTAAGACCAGCTC |
| | P-1819 | GAGCTGGTCTTAGGGATTTG |
| ***LifeAct* GreenGate module** | | aacaGGTCTCtGGCTatgggcgtggccgacctgat |
| | | acaaGGTCTCaCTGAggtggcgaccggtggatc |
| ***mRuby3* GreenGate module** | | aacaGGTCTCaTCAGGAGCAGGGGCGGGTGCCATGGTGTCTAAGGGCGAAGAG |
| | | aacaGGTCTCaGCAGTTACTTGTACAGCTCGTCCATCC |
| ***HSP18.2*-Terminator** | | aacaGGTCTCtCTGCatatgaagatgaagatgaaatatttgg |
| | | aacaGGTCTCaTAGTcttatctttaatcatattccatagtcc |
